# HIV-1 Vpr induces ciTRAN to prevent transcriptional silencing of the provirus

**DOI:** 10.1101/2022.11.04.515166

**Authors:** Vipin Bhardwaj, Aman Singh, Rishikesh Dalavi, Lalchhanhima Ralte, Richard L. Chawngthu, Nachimuthu Senthil Kumar, Nagarjun Vijay, Ajit Chande

## Abstract

The functional relevance of circular RNA (circRNA) expression in HIV-1 infection remains unclear. By developing a customized protocol involving direct RNA nanopore sequencing here, we captured circRNAs in their native state from HIV-1 infected T cells and identified ciTRAN, a **ci**rcRNA modulator of HIV-1 **Tran**scription. We show that HIV-1 infection of monocytic, T cell lines and primary CD4+ T cells induces ciTRAN expression in a Vpr-dependent manner. ciTRAN protein interactome analysis by proximity biotinylation and mass spectrometry identified SRSF-1 as a prominent interactor of the circular RNA. SRSF-1 is known to negatively regulate HIV-1 transcription, which the virus overcomes by a yet unknown mechanism. We demonstrate that HIV-1 Vpr induced ciTRAN sequesters SRSF1 away from the viral transcriptional complex to promote efficient viral transcription. Accordingly, ciTRAN depletion by CRISPR-Cas phenocopied the effects of SRSF1 overexpression and improved SRSF1 association with HIV-1 transcriptional complex. Finally, we show that an SRSF-1-inspired competing peptide can inhibit HIV-1 transcription regardless of ciTRAN induction. The hijacking of a host circRNA thus represents a new facet of primate lentiviruses in overcoming transmission bottlenecks.

## Main

Circular RNAs (circRNAs) are abundant in immune cells, and many of them are regulated during immunological signaling, inflammation, and viral infection (1). However, only a few circRNAs are known to exert regulatory influences in the context of immune function and viral infection(2–5). Notably, the role of circRNAs in various stages of HIV-1 life cycle remains elusive.

CircRNAs are formed by back-splicing events that occur in both coding and non-coding linear transcripts(6, 7). The back-splice junction (BSJ) connects two ends and distinguishes a circRNA from its linear counterpart. By detecting and quantifying the number of distinct BSJ sequences, high-throughput sequencing approaches can be used to characterize cDNA-based circRNA expression in various pathological conditions(8, 9) but are unable to detect circRNA BSJs in their native form. Sequencing platforms, such as Oxford Nanopore Technology (ONT), enable sequencing RNA in the native form, but their potential remains currently limited to capturing polyadenylated transcripts(10).

Circular RNA detection by high-throughput sequencing relies on reverse transcription, which can lead to false circRNA predictions because of the template-switching of the polymerase (11, 12). Moreover, PCR bias can lead to more efficient amplification of some circRNAs, additional events such as trans-splicing and exon repetitions further hamper identification of circRNA in the native form(13). Accordingly, the long-read sequencing technology has been applied to circRNAs only from reverse transcribed products (14–16).

Here, we developed a pipeline for detecting **circ**RNAs by Nanopore **D**irect **R**NA **seq**uencing (circDR-Seq) and identified a host-encoded circRNA (ciTRAN) that is hijacked by HIV-1 to promote viral transcription.

## Results

### Enrichment of circRNAs for direct RNA nanopore sequencing

The ability to reliably identify circRNA depends on the efficiency of the circRNA isolation strategy in a context where linear RNA exceeds by >99% that of circRNA. We sequentially removed non-circRNAs by iterative depletion of the linear RNA species (Figure 1A). Accordingly, total RNA from mock and HIV-1 GFP infected Jurkat E6.1 T cells (Figure S1A) was first subjected to probe-based degradation by RNase H (Figure S1B), which led to more than 95% reduction of 18S and 5S rRNA species without impacting the representative circRNAs (Figure 1B). The remainder linear pool from the ribo-depleted fractions was first polyadenylated using *E. coli* polyA polymerase and subsequently removed using oligo-dT coated magnetic beads (Figure S1C), leading to the depletion of more than 94% and 97% of the abundant linear RNAs from the mock and infected RNA pool, respectively (Figure 1C). The efficiency of RNase R is also limited by the presence of structural RNAs and the availability of diverse substrates, including snRNAs(17). Compared to the RNaseR treatment-based approach, our method (circDRseq) enabled the efficient depletion of G-quadruplexes-containing RNAs (G4-RNA) as well as the small RNA fraction (Figure 1D) while resulting in a 50-86-fold enrichment of representative circRNAs species (Figure 1E).

**Figure 1.**
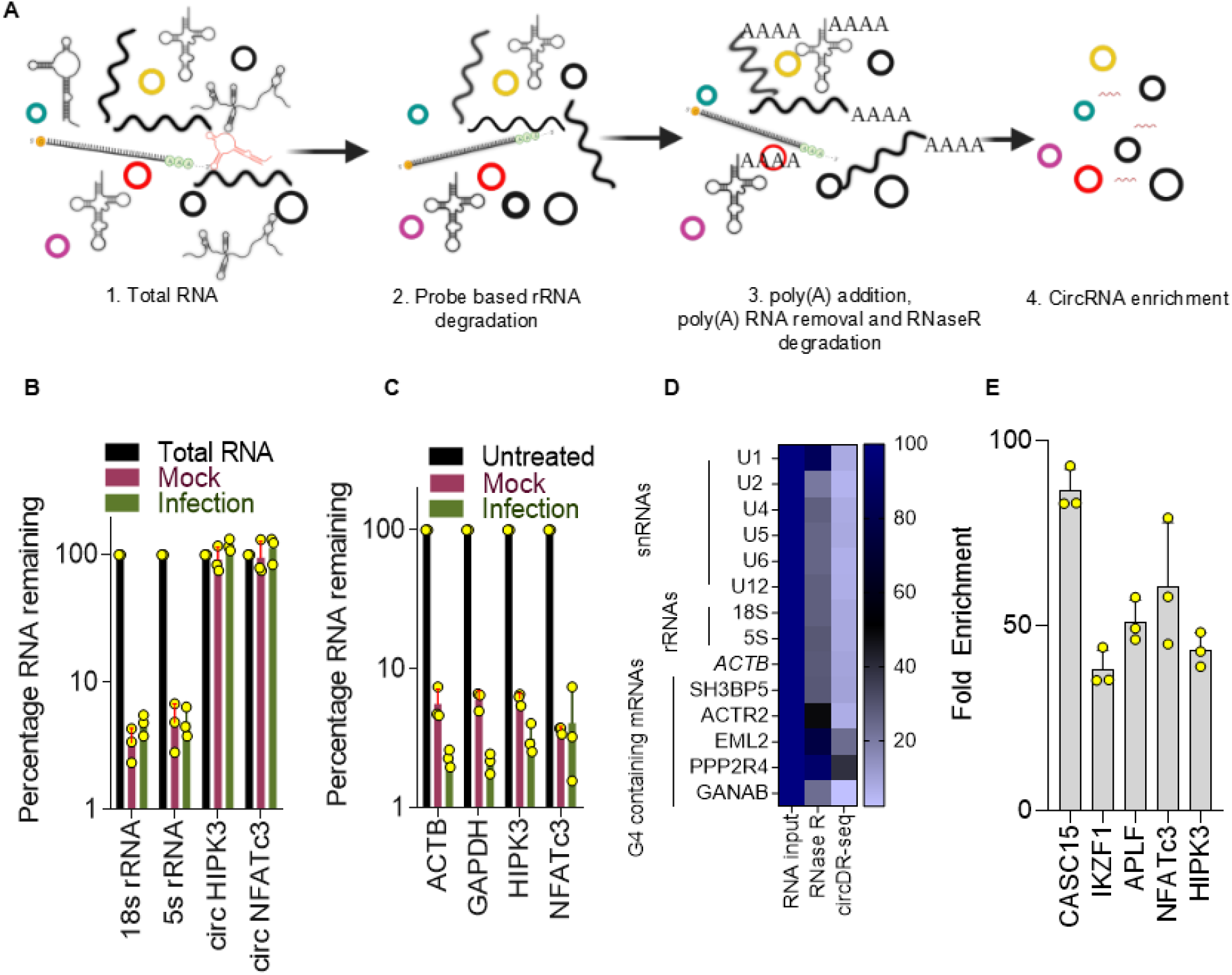
circRNA enrichment by iterative depletion of linear RNA pool. **(A)** Schematics depicting sequential steps to deplete the total RNA of linear RNAs and enrich circRNA fraction. **(B)** RT-qPCR for residual 18S and 5S rRNAs and the indicated circRNAs post-rRNA depletion. The data are normalized to a housekeeping gene *GAPDH*. **(C)** RT-qPCR for residual mRNAs after poly(A) depletion, data normalized to circHIPK3 and cirNFATc3. **(D)** Comparison of RNaseR alone and circDR-seq for assessing levels of subsets of snRNAs, G4 region containing mRNAs, and rRNAs after respective treatments of the mock-treated RNA sample. Data obtained by RT-qPCR are normalized to circHIPK3. **(E)** The extent of circRNA enrichment assessed by RT-qPCR of mock-treated sample for the indicated candidate circRNAs. Data are normalized to GAPDH. n=3;±SD.

### Detection of circRNAs in the native forms

Since circRNAs lack free ends and polyA tail, we first linearised the enriched circRNAs by controlled fragmentation using NEBNext Magnesium RNA fragmentation module (details in the supplementary information). The linear fragments were then polyadenylated to allow direct RNA sequencing for the detection of BSJs by nanopore (Figure 2A). The addition of poly-A tail to the fragmented circular RNA species from Figure 1E was confirmed by amplification from cDNA templates generated by oligo-dT priming (Figure S1D). Next, we generated the library from three replicates, and the tabulated data represents the output from three independent runs (Figure 2B). The representative yeast reads are from an internal standard comprising yeast enolase RNA added in Oxford Nanopore direct RNA sequencing kits. We used NanoPack (a package to visualize and analyze long-read sequencing) to analyze raw data generated from mock and infected samples. The analysis revealed a comparable read length, cumulative yield, basecall quality over time, and average read quality over read length (Figures S2A-H) (details in supplementary information). The next step was to detect BSJs from the sequencing reads obtained. For this, we first extracted circRNAs FASTA sequences from circBase and circAtlas and joined the ends of these sequences (according to the pipeline described in Figure 2C) to obtain the BSJ sequences. Considering the high precision, recall, and F1 score (Figures S3A), we selected a 100 bp library for pblat (Figure 2C; and supplementary information). Using these stringency conditions, we detected 451 and 481 circRNAs with an overlap of 278 circRNAs in the mock-treated sample (Figure 2D). In infected cells with an overlap of 41 circRNA, 76 and 107 circRNA were detected, respectively (Figure 2E). We further expanded our analysis beyond the public databases to identify unannotated circRNAs from our DRS data by generating a virtual exonic library. Interestingly, out of 458 and 86 circRNAs, 151 in mock and 39 in infection were unannotated (Figure S3B). The number of circRNAs was comparatively less in HIV-1 infection, which could result from RNaseL activity (2). We also confirmed two unannotated circRNAs by Sanger sequencing to verify the presence of novel BSJs (Figures S3C and S3D) in our data sets. We found more than 80% of the circRNAs originating from protein-coding genes (Figures S3E and S3F). Comparison of the size distribution and strandedness (Positive or negative DNA strand) corroborated with circBase, eliminating any size and strand-based bias in detection by circDRseq (Figures S3G-K).

**Figure 2.**
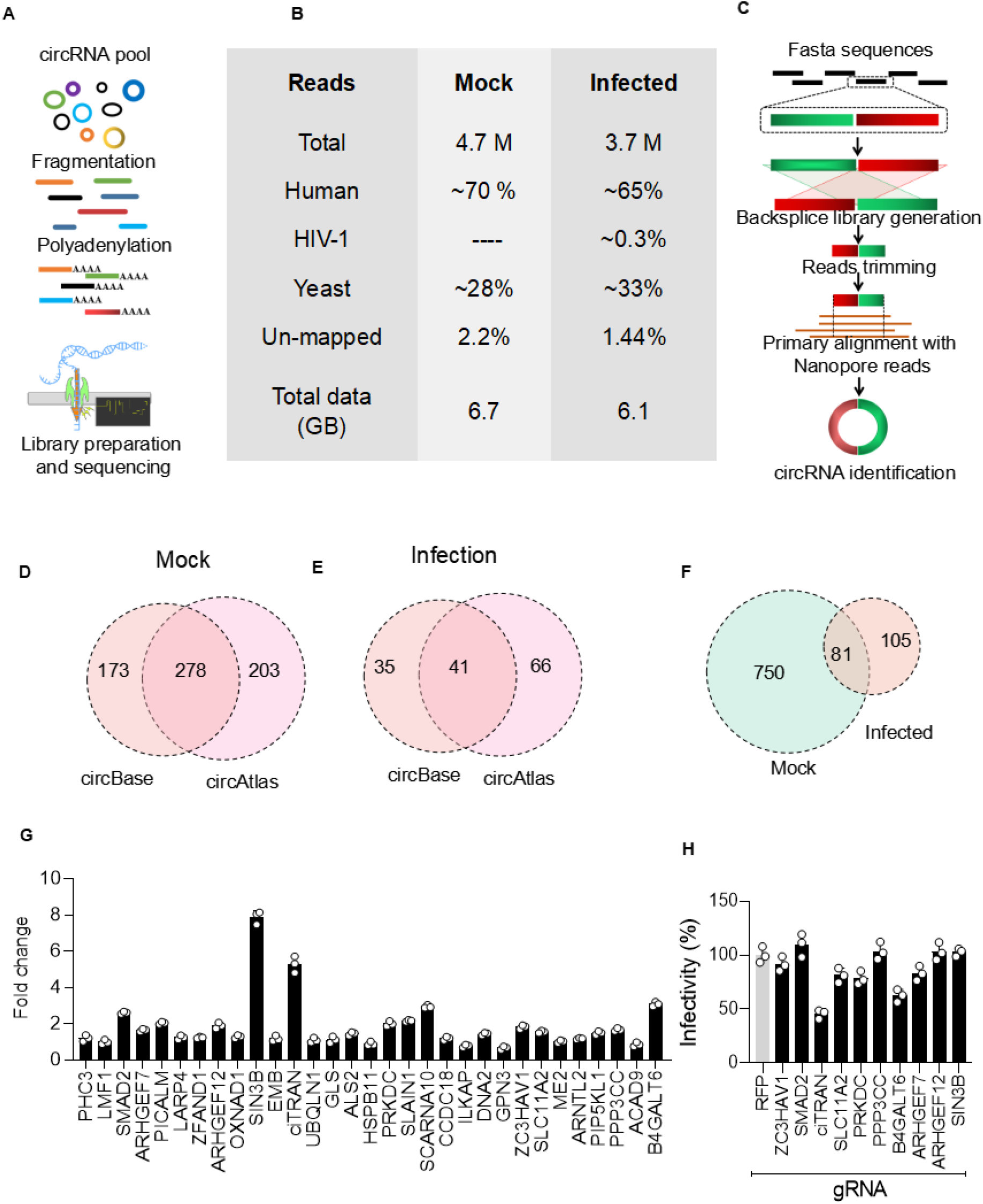
Nanopore DRS reveals infection-induced circRNAs. **(A)** Schematics depicting processing of enriched circRNA pools for Nanopore direct RNA sequencing (Method details in supplementary information). **(B)** Data obtained from 3 flow cells per sample after high precision basecalling. **(C)** Schematics showing a computational pipeline developed for the detection of back-splice junctions using circBase, circAtlas, and virtual exonic circRNA library. **(D,E)** The number of circRNAs obtained after using circBase-specific library in mock (**D**) and infection (**E**). **(F)** The numbers of circRNAs obtained after analysis of virtual exonic circRNA library using a pipeline in C. **(G)** Infection-specific (31 circRNAs) high-confidence circRNA candidates validated using RT-qPCR from HIV-1 infected Jurkat E6.1 cells. Data were normalized to GAPDH. **(H)** Luciferase activity as a function of HIV-1 Luc infection (infection using VSVG pseudotyped NL4-3 Env-R-Luc) from JTAg cells depleted of indicated circRNAs. gRFP served as non-relevant gRNA n=3;±SD.

One of the advantages of direct RNA sequencing by nanopore is that it enables simultaneous detection of m6A modification. RNA m6A modification regulates key functions like controlling RNA splicing, translation, stability, and translocation. Considering circDR-seq can allow simultaneous detection of RNA-associated modifications, we assessed the possibility of detecting m6A containing circRNAs, and their possible coding potential by detecting the presence of m6A and internal ribosome entry site (IRES). Through an integrative analysis (by tools like Nanom6A, CPC2 and IRES finder), we obtained high-confidence protein-coding circRNA candidates. Nanom6A revealed 120 and 21 m6A containing sites that comprised 28 and 7 circRNAs, respectively (Figure S4A,B). Further, randomly selected m6A circRNA candidates were also validated for the presence of the modifications using an N6-Methyladenosine Enrichment Kit (NEB) (Figure S4C,D) that detected the m6A presence on the circRNAs. However, we did not find infection-specific m6A modifications for the candidates tested. Nevertheless, these experiments highlighted the potential of circDRseq for concurrently detecting circRNA modifications. Further, CPC2 and IRES analysis showed an overlap of 98 and 16 circRNAs in mock and infection, respectively (Figures S4E and S4F). Altogether four overlapping circRNAs in mock were detected, which plausibly have the coding potential (Figure S4G). Finally, the robustness in detecting circRNAs was assessed by comparing circDR-seq with the CIRI-long and circNICK-LRS (recently published pipelines) that showed 99.93 and 82.20% efficiency in detection, respectively (Figures S4H-K). Altogether, using circDR-seq, 1017 circRNAs (831 in mock and 186 in infection) candidates were detected in their native forms (Figure 2F), and the possibility of simultaneously capturing RNA modifications was also demonstrated.

Despite low representation, circDRseq enabled the detection of circRNAs from HIV-1-infected T cells in their native states. The next step was to interrogate the functions associated with these RNA species in HIV-1 infection. Out of 41 candidates that were shortlisted based on their common presence in the databases (Figure 2E), five were common to the uninfected pool. The remaining 36, referred to as infection-specific, were selected for further validation. We found a reliable PCR amplification of the desired size for 31 circRNAs. Out of these 31, the expression levels of 10 circRNAs were more than 1.5-fold in the case of infection (Figure 2G). We hypothesized that these upregulated circRNAs could have a role during HIV-1 infection. To find out, we designed Cas13d (CasRx) targeting gRNAs against the BSJs (Figure S5A) for these select candidates (from Figure 2G) and performed a CRISPR/CasRx-based (CasRx or Cas13D is an ssRNA targeting nuclease) knockdown in Jurkat Tag (JTAg) cells. The cells depleted of indicated circRNAs were selected on puromycin, expanded, and knockdown for individual circRNA was confirmed subsequently using RT-qPCR (Figure S5B; circRNA-specific primers tabulated in Supplementary Table S1). Interestingly, the HIV-1 infection in circRNA-depleted target cells by luciferase-encoding HIV-1 (HIV-Luc) revealed that ciTRAN (55%) and, to a lesser extent (37.5%) circB4GALT6 reduced the luciferase expression from the virus scored as a function of infection (Figure 2H). We chose to examine the impact of ciTRAN expression on HIV-1 lifecycle because of its ability to reduce infectivity.

### ciTRAN relieves a post-integration block to viral gene expression

To investigate the step of the virus life cycle impacted by ciTRAN, the target cells that either expressed a non-relevant gRNA (Luciferase gRNA) or ciTRAN-specific gRNA were used (Figure S5C). The subcellular fractions from the HIV-1 GFP infected cells were used to capture reverse transcription products, integration events, and proviral transcription. In conditions where ciTRAN, but not the linear counterpart, was depleted (Figure S5D), our experiments suggested that depletion of ciTRAN did not impact reverse transcription from the viral genomic RNA (Figure 3A), nor was the copy number of the integrated provirus changed (Figure 3B). On the contrary, cytoplasmic viral RNA was reduced, and nuclear run-on assay indicated a 2.3-fold reduction in transcription of HIV-1 RNA in ciTRAN depleted conditions (Figures 3C and 3D). The fewer transcripts consequentially reduced intracellular viral capsid as revealed by western blotting (Figure 3E), pointing to an effect of ciTRAN at a post-integration step.

**Figure 3.**
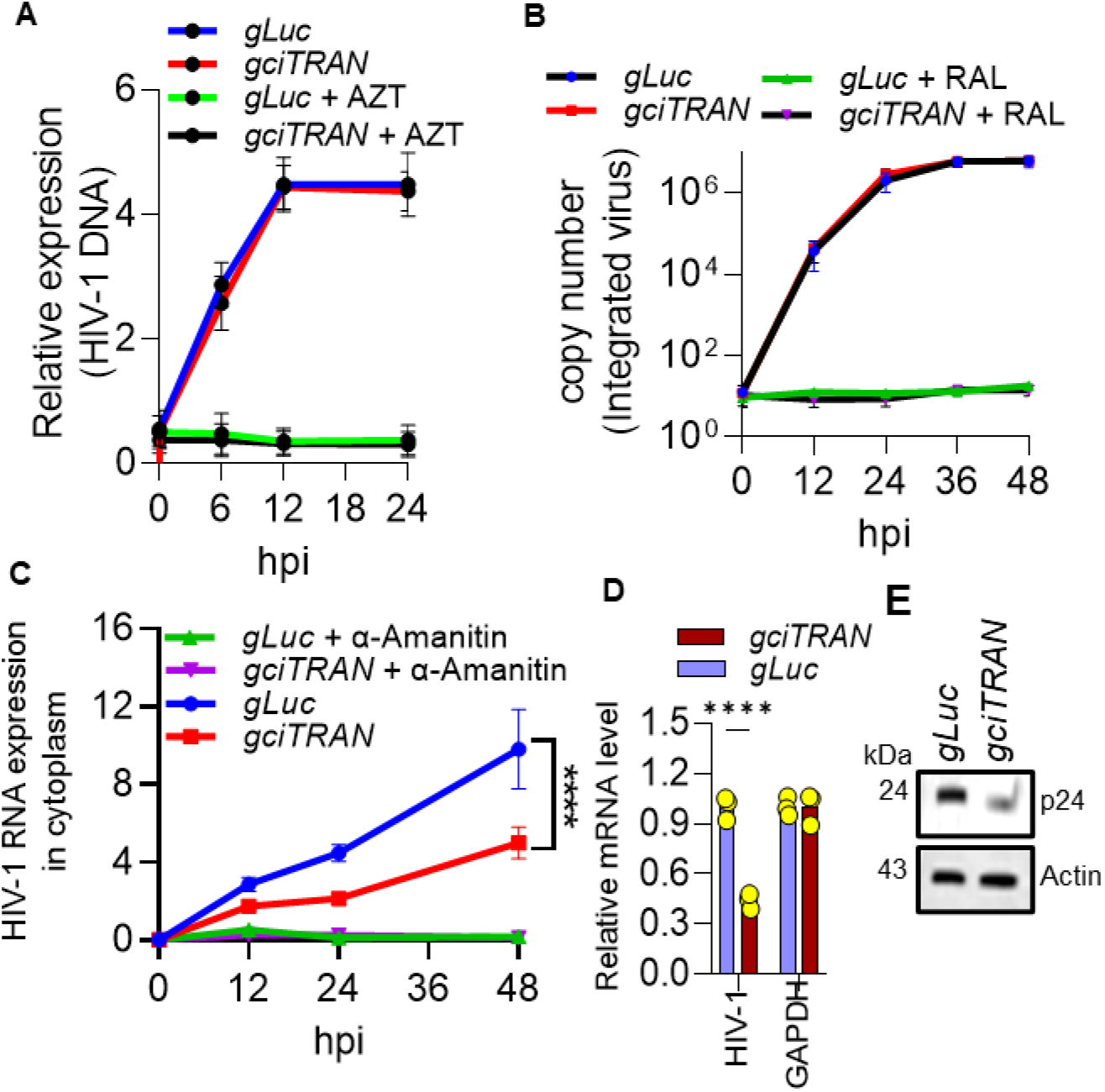
A post-integration event is targeted by ciTRAN. Effects of ciTRAN knockdown during HIV-1-GFP infection on: **(A)** reverse transcription (RT) products as assessed by measuring cytoplasmic HIV-1 DNA (AZT served as a control), **(B)** HIV-1 genomic DNA integration (Raltegravir (RAL) served as a control), and **(C)** cytoplasmic HIV-1 RNA expressed from the provirus (α-Amanitin served as a control). **(D)** Nuclear transcription from an integrated provirus in ciTRAN depleted condition assessed by nuclear run-on assay, GAPDH served as control. gRNA to luciferase served as a non-relevant gRNA **(E)** Immunoblot indicating intracellular viral capsid levels; Actin served as a loading control. n=3;±SD.

### ciTRAN binds to a negative regulator of HIV transcription

CircRNAs can regulate parental gene expression since they can interface with DNA or RNA at the sequence level and establish specific interactions with proteins (18). Furthermore, circRNAs can also directly establish interactions with chromatin or regulate parental gene expression by binding to protein mediators of transcription (19, 20). ciTRAN is encoded by *SMARCA5*, and the circular RNA species is predominantly nuclear-localized (28). Our experiments indicated that ciTRAN depletion negatively impacts transcription from the provirus, so we wondered if ciTRAN targets a host factor to modulate such events. To address this, we employed the CARPID (CRISPR-assisted RNA–protein interaction detection method) (21) for isolating the ciTRAN-interacting proteins. Specific guides were generated that installed a catalytically dead CasRx fused with biotin ligase on the BSJ of ciTRAN. This is followed by a pulse labeling of biotin and subsequent enrichment of the biotinylated pool by streptavidin affinity matrix (Figure S5E). Isolation of differentially enriched bands and identity by mass-spectrometry revealed SRSF1 as a top candidate interactor of ciTRAN (Figure 4A and Figure S5F). PAR-CLIP (photoactivatable ribonucleoside-enhanced crosslinking and immunoprecipitation) (Figure 4B) from the lysates of infected T cells, reciprocally, showed 6-fold enrichment of ciTRAN with SRSF1-specific antibody over IgG control, further confirming the specificity of interaction (Figure 4C). We used HIV-1 Tat, an RNA-binding protein, as non-relevant control that did not co-enrich ciTRAN. This experiment also ruled out the possibility of ciTRAN sequestering Tat and concurrent modulation of the viral transcription (Figure 4D). We, therefore, next examined whether the effect of ciTRAN depletion on HIV-1 gene expression resulted from the loss of interactions between the circRNA and SRSF1. JTAg ectopically expressing haemagglutinin (HA) tagged SRSF1 phenocopied the effects of ciTRAN downregulation by reducing the viral infectivity (Figures 4F). Moreover, SRSF1 expression in the produer cells (JTAg) also reduced the number of progeny virions produced and as a consequence less number of target cells (TZM-GFP) were infected, suggesting an impact on the biogenesis and onward transmission (Figure 4G). Further, regulating the ciTRAN and SRSF1 levels in non-T cells (HEK293T) cells during virus production also influenced the number of infection events produced by the progeny virions, suggesting that the phenotype is not cell type-specific (Figure 4H).

**Figure 4.**
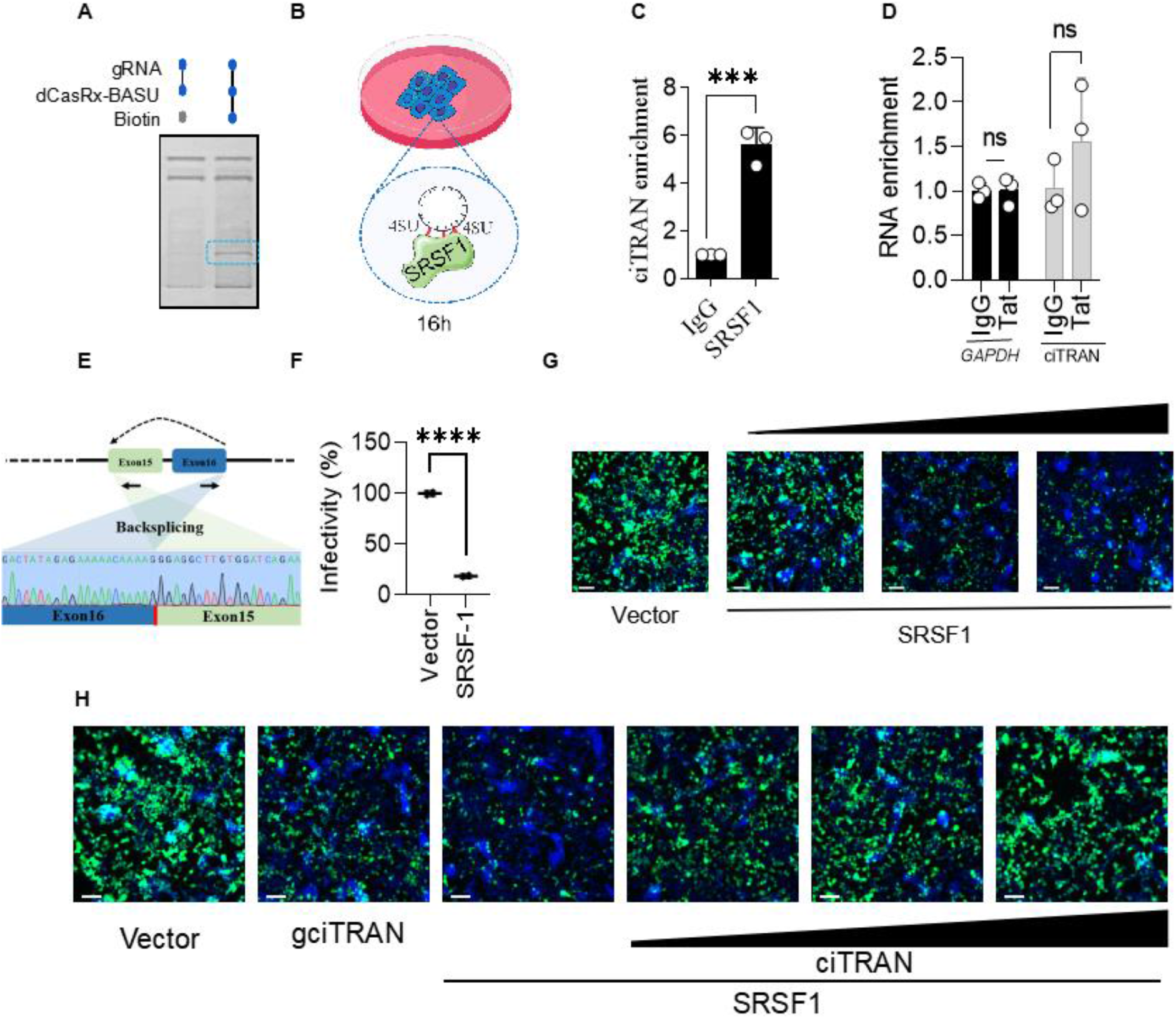
ciTRAN interacts with a negative regulator of HIV-1 infectivity. **(A)** Biotinylated protein fractions obtained by ciTRAN mediated dCas13d-BASU based proximity labelling visualized by silver staining. The differentially resolved band is highlighted. **(B)** Schematics for SRSF1 PAR-CLIP to validate ciTRAN association. **(C)** SRSF1 PAR-CLIP followed by qPCR using ciTRAN and GAPDH specific primers shows enrichment for ciTRAN; n=3;±SD. **(D)** HIV-1 Tat PAR-CLIP followed by qPCR using ciTRAN and GAPDH specific primers shows no difference in enrichment; n=3;±SD. **(E)** Backsplicing junction of enriched ciTRAN RNA during SRSF-1 PAR-CLIP from Figure 4C was validated using Sanger sequencing. **(F)** Effect of SRSF1 ectopic expression on HIV-1 infectivity in JTAg cells; n=3;±SD. JTAg cells were electroporated with pCDNA 3.1(-) and pCDNA 3.1(-) SRSF-1 HA and then were infected using VSVG pseudotyped HIV-1-Luc (NL4-3 E-R-Luc). **(G)** HIV-1 produced by electroporation from JTAg cells along with vector and increasing concentration of SRSF1 (pCDNA 3.1 SRSF1-HA, 500 ng, 1000 ng and 2000 ng) and infectivity in TZM-GFP target cells, representative images (Scale 100 μm). **(H)** Effect of ciTRAN knockdown, ciTRAN ectopic expression(pCDNA 3.1(+) circMini ciTRAN, 100ng,150ng and 200ng) and SRSF1 expression (50ng) on HIV-1 infectivity. HIV-1 produced from HEK293T in the presence of either vector, gciTRAN (ciTRAN knockdown cells), synthetic ciTRAN expressing vector (pCDNA 3.1(+) circMini ciTRAN), or SRSF1 expression vector (pCDNA 3.1(-) SRSF-1 HA). Representative images of TZM-GFP target cells (Scale 100μm).

### ciTRAN obstructs LTR accessibility of SRSF1 to promote infectivity

To obtain detailed insights into the ciTRAN-SRSF1 axis and its contribution to the formation of the functional transcriptional complex on HIV-1 promoter, we examined SRSF1, RNAPII, and Tat occupancy on HIV-1 promoter (TAR (trans-activating region) locus) by chromatin immunoprecipitation (ChIP). ChIP experiments indicated a strong interaction of SRSF1 with the TAR locus (Figure 5A and Figure S5H). Strikingly, this association of SRSF1 with the TAR locus was diminished by overexpression of ciTRAN from a synthetic construct, and the ectopic expression of SRSF-HA restored it to 70%, suggesting a competing scenario (Figure 5A). We also checked the association of RNAPII on the HIV-1 promoter in these identical conditions and found that increased SRSF1 association excludes Pol-II from the viral promoter and that ciTRAN restores the polymerase association for effective transcription (Figure 5B and Figure S5I). SRSF1 is a constitutively expressed protein that competes with and replaces HIV-1 Tat from the transcription complex (22). In agreement, the reduced signal for Tat ChIP was suggestive of Tat displacement by SRSF1 from the transcribing locus (Figure 5C and Figure S5J). Altogether these experiments indicated that ciTRAN competes for SRSF1 sequestration to reinstate Tat interaction.

**Figure 5.**
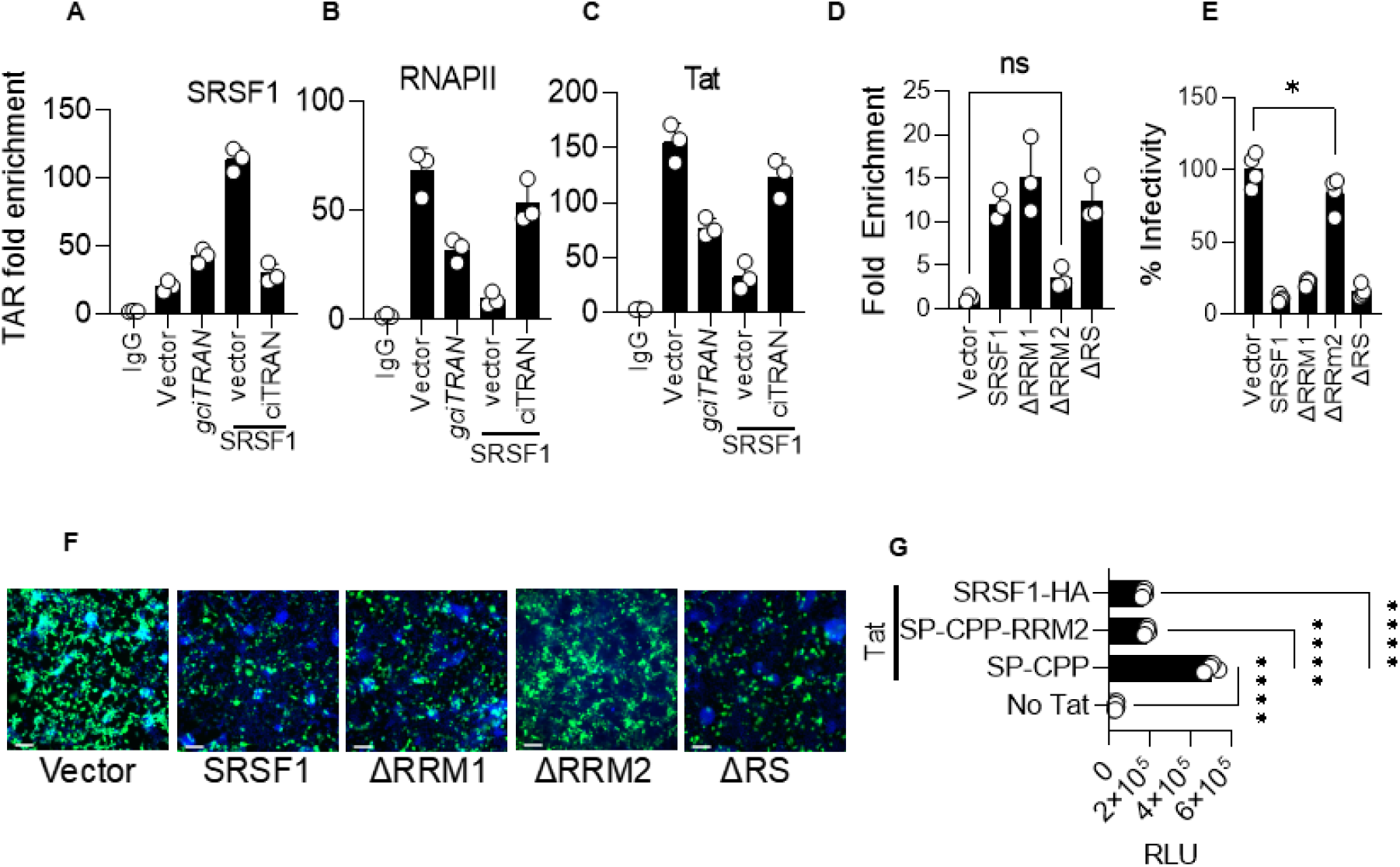
SRSF-1 sequestration by ciTRAN promotes HIV-1 transcription. (A-C), Chromatin immunoprecipitation analysis from JTAg cells and quantification of TAR locus enrichment by RT-qPCR for SRSF-1 (A), RNAPII (B), and HIV-1 Tat (C); n=3;±SD. HIV-1 (NL4-3 Env-Nef-) vector was electroporated in the JTAg cells along with vector, gciTRAN, SRSF-1 with empty vector pcDNA 3.1(+) circMini and pCDNA3.1(+) circMini encoding ciTRAN. Line highlighting SRSF-1 shows SRSF1 expression along with empty vector and ciTRAN ectopic expression. The fold enrichment method was used to represent the data. (D) Effect of SRSF-1 deletion mutants on ciTRAN interaction assessed by PAR-CLIP. SRSF-1 deletion mutants were ectopically expressed using pCDNA 3.(-) HA construct in JTAg cells using electroporation and PAR-CLIP was performed by pulling down mutants using anti-HA antibody. (E,F) Effects of indicated SRSF1 deletion mutants on single-cycle HIV-1 infectivity (n=4;±SD) and representative images (F) from TZM-GFP cells (Scale 100μm). HIV was produced from JTAg cells by electroporation, and the infection was given in TZM-GFP target cells and scored by counting green cells. (G) Effect of CPP-tagged RRM2 on HIV-1 LTR activity assessed by LTR-driven luciferase activity. PGL3 harboring NL4-3 LTR upstream of firefly luciferase was co-transfected in HEK293T cells along with HIV-1 Tat and SP-CPP or SP-CPP-RRM2 and pCMV Renilla-luc. Firefly luciferase readings were assessed as measure of HIV-1 LTR activity in presence of RRM2 and empty vector. Data was normalized with Renilla luciferase. SRSF-1 taken as positive control for LTR inhibition.

### RRM2-inspired peptides can prevent ciTRAN action

SRSF1 contains two RNA recognition motifs (RRMs) and a carboxy-terminal rich in arginine/serine-rich (RS)(22, 23). We next examined the molecular features of SRSF1 required for ciTRAN interaction and the consequential effects on viral transcription in the absence of those. Employing a series of deletion clones (Figure S5G and S5K), using PAR-CLIP, we found that when RRM2 was deleted, there was no enrichment of ciTRAN (Figure 5D). The loss of ciTRAN interaction in ΔRRM2 mutant is also consistent with its ability to promote viral infectivity (Figures 5E and 5F).

Next, we sought to explore SRSF1protein domains as viral gene expression inhibitors in the context of ciTRAN expression. Accordingly, we generated a cell-penetrating peptide (CPP) tagged RRM2 and showed that it can inhibit virus gene expression, suggesting the possibility of therapeutically targeting the ciTRAN-SRSF1 axis (Figure 5G). Collectively, these experiments provided molecular determinants of SRSF1 required for ciTRAN recognition, mechanistic insights into the ciTRAN-SRSF1 axis, and therapeutic targeting possibility.

### HIV-1 Vpr induces ciTRAN expression

Having established the ciTRAN-SRSF1 axis in HIV-1 transcriptional regulation and its virological effects, we next investigated the viral side of the equation in regulating the phenotype. Expression of ciTRAN was observed in natural infections (n=15) ranging from 4-13-fold (Figure 6A). Additionally, when PBMCs and purified primary CD4+ cells isolated from healthy donors were infected with HIV-1 (Figures S6A, S6B,S6C), we observed up to 10-fold induction of ciTRAN (Figures 6B and 6C). The ciTRAN upregulation was only limited to HIV-1, and MLV could not induce it, as revealed by ectopic expression of HIV-1 and MLV constructs in JTAg cells (Figure 6D). These results implied the relevance of ciTRAN expression in HIV-1 infected primary cells and cell lines and provoked us to surmise the involvement of virally encoded proteins in stimulating the ciTRAN expression. To find out, we assessed ciTRAN levels by transfecting the virus genomes that could not produce indicated viral accessory/regulatory proteins and enzymes. ciTRAN levels were unchanged in almost all conditions except when the virus lacked Vpr; the ciTRAN levels were equivalent to the empty vector-transfected pool when the virus lacked Vpr (Figure 6E). Further, the infection of primary CD4^+^ cells with Vpr +/−HIV-1 Luc viruses confirmed the involvement of Vpr accessory protein in the induction of ciTRAN (Figures 6F, 6G and 6H). We next investigated whether Vpr requires help from other viral proteins to effect ciTRAN induction. Representative alleles expressed from pCDNA 3.1(-) from NL4-3, transmitted founder viruses CH040 and REJO indicated that it is a conserved feature associated with HIV-1 encoded Vpr. In contrast, Vpx cannot induce ciTRAN expression (Figure 7A). Vpr sufficient HIV-1 is associated with higher virion infectivity in T cells compared to HIV-1 deficient in Vpr. We show that ciTRAN contributes to this infectivity enhancement (Figures 7B and S6D. In agreement with previous reports (24, 25), we found that HIV-1 Vpr enhances the viral replication in human macrophages via various mechanisms (Figure 7C). Further, virion-encapsidated Vpr also has been implicated in the infection of macrophages. Therefore, we next asked if virion-packaged Vpr is competent to increase cellular levels of ciTRAN. THP-1 monocytes challenged with HIV-1 Luc showed more than 10-fold induction of ciTRAN, and that virion-encapsidated Vpr (during the production, Vpr is packaged through expression *in trans* using pEGFP-C2) was sufficient to induce the circRNA (Figures 7D and 7E). Altogether, these results indicated that HIV-1 Vpr is required and sufficient to induce the expression of ciTRAN in monocytes and T cells.

**Figure 6.**
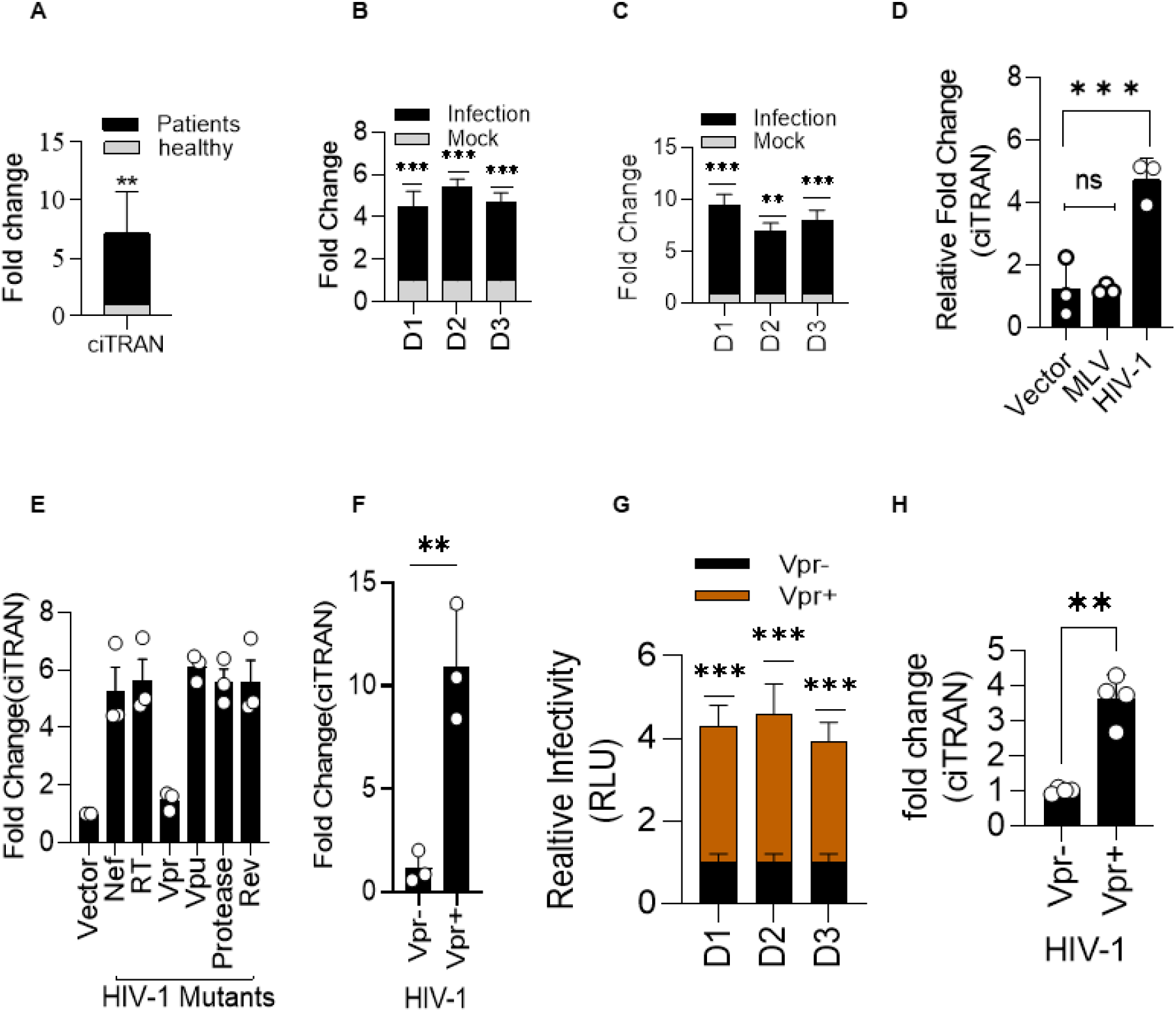
ciTRAN is induced by Vpr. (**A**) ciTRAN level analysed using RT-qPCR in total RNAs isolated from whole blood of healthy donors (n=5) and HIV infected individuals (n=15). **(B,C)** Effect HIV-1 infection on ciTRAN levels in the PBMCs (**B**) and CD4+ cells (**C**) isolated from three healthy donors (D1-3). HIV-1 GFP was used for infecting PBMCs and CD4+ cells for 48 h. RNA from infected and mock treated cells were used for ciTRAN level estimation using qRT-PCR; n=3 ±SD. **(D)** Effect of MLV and HIV-1 on ciTRAN induction in JTAg cells; n=3±SD. MLV encoding plasmid NCA wt zsGreen and NLBN zsGreen was electroporated in JTAg cells for 48h to analyze the ciTRAN induction. RNA from these cells were used for qRT-PCR analysis for ciTRAN analysis. Data was normalized using house keeping gene, GAPDH. **(E)** Effect of viral accessory proteins on ciTRAN induction estimated using RT-qPCR after electroporation of JTAg cells with HIV-1 plasmids lacking indicated viral genes(described in resource table in supplementary information); n=3 ±SD. **(F)** Effect of Vpr on ciTRAN levels in CD4+ cells assessed by RT-qPCR after HIV-1 Luc Vpr (+/−) virus infection; n=3 ±SD. HIV-1 produced with and without Vpr from HEK293T was used to infect CD4+ cells. (**G**) HIV-1 Luc Vpr+/−infectivity in CD4+ primary T cells. (**H**) ciTRAN level revealed by RT-qPCR in primary CD4+ cells infected with HIV-1 Vpr+/-.

**Figure 7.**
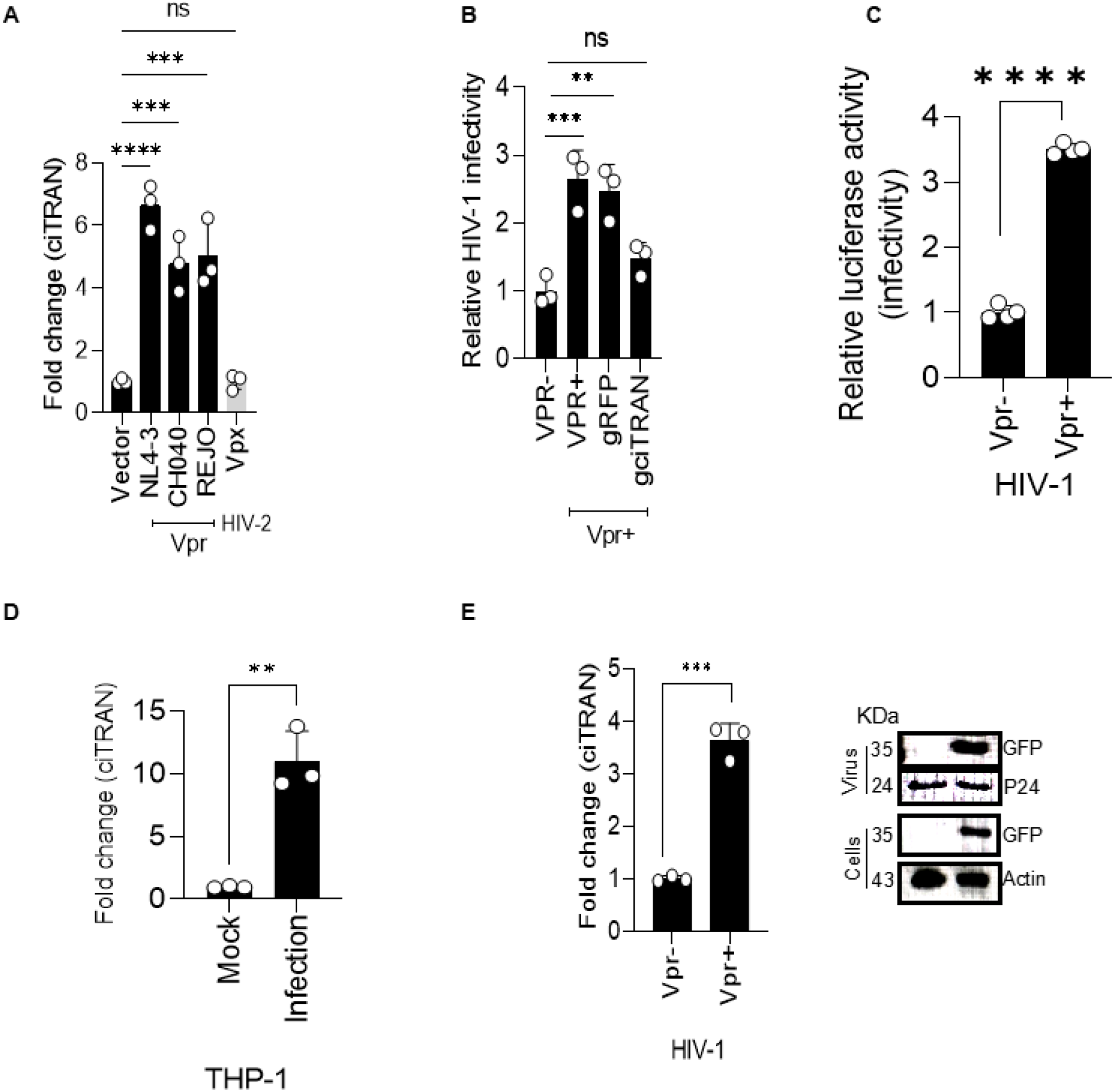
ciTRAN induction by Vpr is conserved and is not T cell-specific. **(A)** Effect of ectopic expression of HIV-1 Vpr alleles and Vpx from HIV-2 on ciTRAN induction in JTAg T cells, GAPDH served as a reference for normalization; n=3 ±SD. All the Vpr alleles were ectopically expressed through a pcDNA backbone (described in supplementary information) **(B)** Effect of ciTRAN knockdown on Vpr+/−HIV-1 Luc infectivity in JTAg cells. ciTRAN knockdown JTAg T cells made using Cas13d were challenged with HIV-1-Luc with and without Vpr. Infectivity was scored as relative luciferase units normalized to Bradford protein quantification from cell lysate to nullify the cell number effect. **(C)** Vpr+/−HIV-1 Luc infectivity as a measure of relative luciferase units in THP-1 monocytes normalized to Bradford protein quantification from cell lysate; n=4 ±SD. **(D)** ciTRAN levels in THP-1 monocytes upon HIV-1 infection; n=3 ±SD. THP-1 cells were infected with HIV-1 Gfp. **(E)** Effect of virus encapsidated Vpr on ciTRAN level in THP-1 monocyte cells (left panel) and immunoblotting showing Vpr expression in viruses and in target THP-1 cells (right panel). HIV-1 Luc (NL4-3 E-R-Luc) produced from HEK293T in presence of empty vector pEGFP or pEGFP C2 Vpr, and the infection was given in THP-1 cells. Vpr packaging inside the particles was confirmed using immunoblotting using anti-GFP antibody.

### ciTRAN-SRSF1 axis is functionally conserved

Identification of a well-preserved SRSF1 binding site on LTR from various HIV-1 isolates (Figure S6E) reinforces the conservation of ciTRAN-SRSF1 axis in regulating the viral transcription. We, therefore, experimentally checked if ciTRAN regulates LTR transcription for transmitted founder viruses (TFV) in addition to LTRs from other clades. To experimentally assess ciTRANs activity on LTR-driven expression, we generated PGL3-Luc-based minigene LTR reporters (Figure S6F) and performed luciferase assays in HEK293T cells by co-transfecting LTR minigenes along with Cas13Rx and gRNA against ciTRAN or non-targeting gRNA. To confirm the effect of ciTRAN depletion on reporter expression. In conditions where ciTRAN was depleted (Figure S6G), the concomitant reduction in luciferase expression from viral LTR was observed regardless of the origin of the LTR (Figure 8A). Additionally, the SRSF1 ectopic expression (pCDNA 3.1(-) SRSF-1 HA ectopic expression in target cells) markedly reduced luciferase expression. However, ciTRAN expression from synthetic construct (pCDNA 3.1(+) circMini ciTRAN) rescued this transcriptional defect for all the LTRs in these conditions (Figure 8B), signifying a conserved ciTRAN-SRSF1 axis in the regulation of HIV-1 transcription.

**Figure 8.**
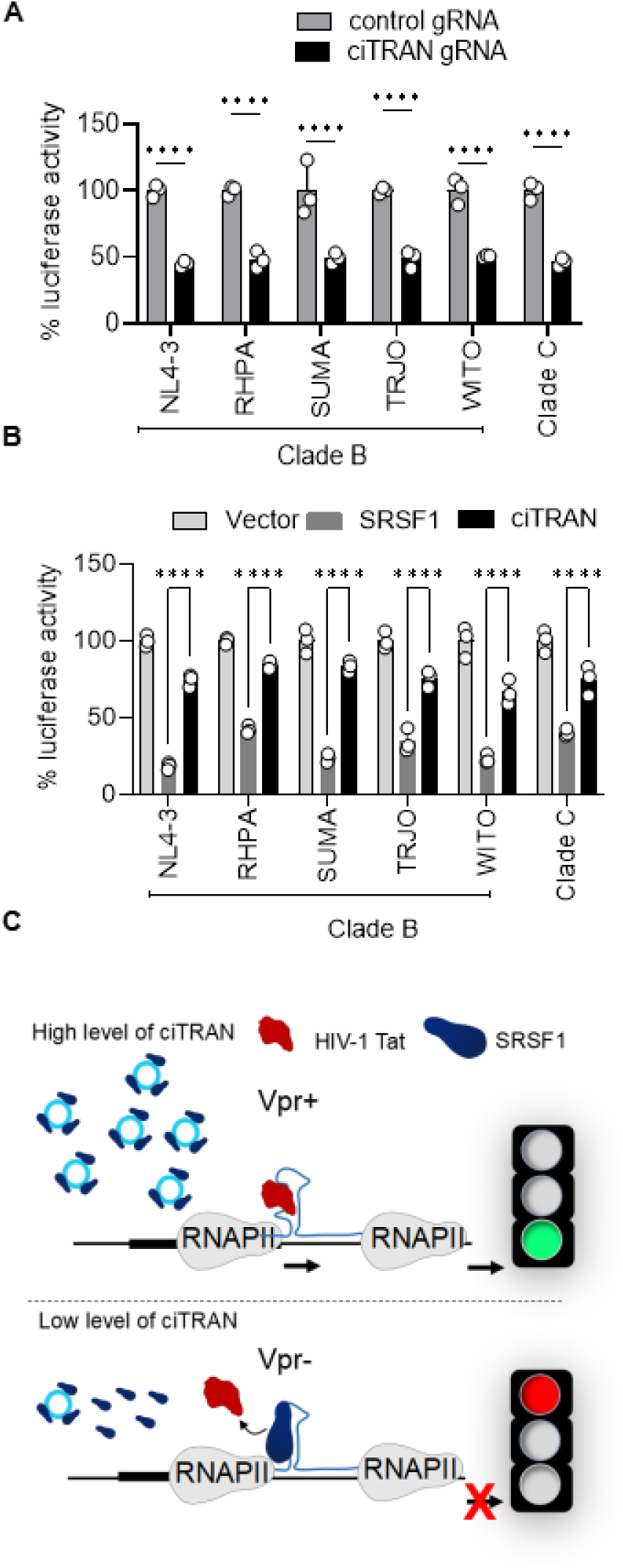
ciTRAN-SRSF axis is functionally conserved. **(A)** Effect of ciTRAN knockdown in HEK293T cells on the reporter activity driven by LTRs from transmitted founder viruses and different clades of HIV-1; n=3 ±SD. **(B)** Effect of ciTRAN and SRSF1 ectopic expression in HEK293T on reporter activity driven by LTRs from transmitted founder viruses and different clades of HIV-1; n=3 ±SD. **(C)** Schematics showing the effect of Vpr-induced ciTRAN on HIV-1 transcription.

SRSF1 expression remained unaltered in the infection (Figure S6H), indicating that a constitutively expressed protein critical for cell survival is a bottleneck for HIV-1 transmission. Our results suggest that Vpr overcomes this bottleneck by hijacking ciTRAN, which sequesters SRSF1 away and reinstates Tat for efficient virus transcription (Figure 8C).

## Discussion

Low expression of circRNAs imposes restrictions on our ability to capture them in the native form, more so from HIV-infected cells due to the over-representation of viral reads, expression of reverse transcriptase, and suppression of host transcription in general(26, 27). By substantially reducing the linear RNA content, we demonstrated that direct RNA sequencing could be used for detecting circRNA BSJs, thus expanding the utility of ONT for not only capturing circRNAs in the native form but also associated modifications. Moreover, although ONT DRS sequencing results in low-read numbers (1-2 million reads/flow cell) compared to hundreds of millions of reads that can be obtained using cDNA-based Illumina platforms, we showed that enrichment using successive linear RNA removal steps could, to a great extent, circumvent the need to obtain a higher number of reads without compromising the data quality. A multistep purification procedure and BSJ detection method might introduce a bias in the circRNAs that are being identified. Therefore, we performed a systematic and extensive validation of circDR-seq: using different databases, simulated datasets, and comparative assessment with recently published protocols (CIRI-long and circNICK-LRS), size distribution, experimental and functional validation, which exemplify the reliability of our approach in detecting bonafide circRNAs. Further, circDR-seq enabled the detection of circRNAs of sizes ranging from as small as 100 bases to more than 5000 bases, which is currently considered a limitation by cDNA-based circRNA nanopore sequencing approaches (Rahimi, 2021, Zhang, 2021). Further, our identification of unannotated BSJs in the native form and further validation by Sanger sequencing exemplify the utility of circDRseq in cataloging novel circRNAs. circDR-seq can potentially be extended to capturing circRNAs of various sizes and understanding their contributions to other pathophysiologies.

HIV-1 infection-associated circRNAs were reliably captured, and we further demonstrated a functional role for a circRNA ciTRAN in intersecting a molecular pathway involved in HIV-1 transcription regulation. For this, complementary approaches involving cell lines, primary cells, conventional RNA interference, and contemporary RNA manipulating tools were employed, ensuring that the phenotype is not affected by the cognate linear RNA, cell types, or a method of perturbation. We acknowledge that the extent of circRNA depletion by CRISPR was only up to 50% for some circRNAs, and this we purview as a limitation because other than BSJs, we cannot specify the regions for gRNA targeting without impacting the linear counterparts. Thus, methods that achieve higher knockdown efficiency may reveal additional candidate circRNAs and their relevance for infection.

SRSF1 and HIV-1 Tat compete for binding onto an overlapping sequence in the TAR(22, 23). We demonstrated that RRM2 is required and alone is as effective as full-length protein in obstructing Tat binding with TAR. SRSF1, which is constitutively expressed, can out-compete Tat binding in the early phase of viral transcription due to the low level of Tat, thus directly affecting HIV-1 trans-activation (23). We show that ciTRAN intercepts SRSF1’s ability to displace Tat from the transcriptional complex by sequestering it away, enabling efficient viral transcription. Remarkably, the binding of ciTRAN to SRSF1 is also RRM2-dependent, and correspondingly, SRSF1 lacking RRM2 enhances the HIV-1 infectivity.

Therefore, it would be interesting to define the propensity of RRM2 towards ciTRAN binding by structural studies. Furthermore, we show the feasibility of targeting the RRM2-ciTRAN axis by generating a synthetic CPP-tagged RRM2-mimic which we showed antagonizes virion-induced ciTRAN. Future studies can refine this approach by developing small molecule mimics or peptides for inhibiting HIV-1 transcription, an event crucial for virion biogenesis and propagation.

HIV-1 accessory protein Vpr is crucial in the early stage of the viral life cycle in myeloid and lymphoid cells (29–31). Vpr interacts with various host cellular proteins affecting multiple key activities such as nuclear import of pre-integration complex (PIC), apoptosis, G2/M cell cycle arrest, antagonizing HUSH mediated epigenetic silencing, and enhancing viral trans-activation (30, 32, 33). The well-known functions of Vpr, like G2 arrest, and counteraction of HUSH does not explain its effect on transcription and requirement during early phase, suggestive of additional functions of this accessory protein in natural infection(34, 35). Our experimental demonstration that HIV-1 lacking Vpr fails to induce ciTRAN, and virion encapsidated Vpr is sufficient to induce ciTRAN for relieving the SRSF1-mediated transcriptional block provides mechanistic insights into this long-sought Vpr function. Our results are also consistent with encapsidated Vpr’s ability to modulate HIV-1 replication in the early stage of infection(36). Like other auxiliary proteins, Vpr is known to be conserved within different HIV-1 clades. Upon investigation, we found that different Vprs, but not HIV-2 Vpx, can induce the ciTRAN, suggesting a conserved function of this accessory protein. However, how Vpr augments ciTRAN is yet unclear and requires further investigations. Vpr mutational studies could be one way to understand ciTRAN induction and associated enhanced viral transcription.

In sum, hijacking of a host circRNA to avoid RBP-mediated transcriptional silencing offers new insights into the transcriptional regulation of HIV-1 and offers new possibilities for therapeutic targeting.

## Acknowledgments

This work was supported by the DBT/Wellcome-Trust India Alliance fellowship (IA/I/18/2/504006) awarded to AC. VB, RD and AS is supported by a fellowship from CSIR and DBT, respectively. The authors are grateful to Massimo Pizzato, and the NIH AIDS reagent program for reagents and cell lines. All lab members especially Pavitra Ramdas are acknowledged for helpful discussions, various reagents, and technical help.

## Author contributions

Resources: AC, NV, LR, RC, NSK. Data curation: VB, AS, AC. Software: AC, VB, AS, NV. Formal analysis: VB, AS, LR, RC, NSK, NV, AC. Validation: VB, AS, RD, AC. Investigation: VB, AS, AC Methodology: VB, AS, RD, LR, RC, NSK, NV, AC. Conceptualization: AC, VB. Supervision: AC. Funding acquisition: AC.

## Competing interests

The authors declare no competing financial interests.

## Data and materials availability

The Direct RNA sequencing data generated in this study is deposited to European Nucleotide Archive (ENA) with the accession number **PRJEB52141**. The code used for analyzing the data in this study is available on GitHub (https://github.com/aman21392/circDR-seq). The list of reagents, antibodies, and plasmids is given in the supplementary Table1. Oligos used and the raw data related to circular RNAs after various analysis is available in supplementary tables (Tables S1-S7).

## Ethics statement

The Institute Ethics Committee of IISERB approved the PBMC collection from healthy donors with due written consent from the donors. The HIV-positive patient’s sample collection was approved by the Institutional Human Ethics Committee of Mizoram University (MZU//IHEC/2019/007).

## Methods

### Cell culture

Jurkat E6.1 (ATCC), THP-1 (ATCC) and Jurkat Tag (JTAg) (37) cell lines were cultured in RPMI (Biowest) supplemented with 10% fetal bovine serum (Certified, heat-inactivated serum from Gibco; Cat. No. 10082147, lot no. 2097440). PBMCs and CD4+ primary T cells were first activated using PHA (5μg/mL) and IL2 (50 IU) and maintained in RPMI. Primary cells post-activation were expanded in IL2 containing RPMI^39^. HEK293T (ECACC), TZM-GFP (37) cells were maintained in 10 % FBS containing Dulbecco’s modified Eagle medium with 2mM L-glutamine. The cells were maintained in humidified 5% CO2 incubator at 37°C. Cell monolayers were grown at a split ratio of 1:10 by treating with trypsin (0.25%) and 1 mM EDTA (Invitrogen, Thermo Scientific, USA). Detailed description of the reagents is provided in Table S1.

### Virus production and infection

For Target cell infection, the virus was produced from HEK293T producer cells using the calcium phosphate method by co-transfecting 7 μg NLBN zsGreen Env defective and Nef defective (HIV-1 GFP) and 1 μg pMD2.G encoding VSVG glycoprotein and was limited to single-cycle replication. NLBN zsGreen was a kind gift from Prof. Massimo Pizzato which is slightly modified from NL4-3 Env-Nef-described in Rosa et al. Nature 2015 where env region is deleted and zsGreen is cloned in place of Nef. The virus-containing culture supernatant was collected after 48h post-transfection, centrifuged at 300xg for 5 min, and then filtered using a syringe filter of 0.22μ. Virus was quantified using SGPERT assay(38) and multiplicity of infection (MOI) was calculated by infecting HEK293T cells. Jurkat E6.1 cells were infected at MOI=2, whereas primary CD4+ T cells and CD14+ T cells were infected at MOI=5. For Nanopore sequencing, E6.1 cells were infected in a T75 flask (Eppendorf, Germany) for 48h and processed for RNA isolation. Infectivity was quantified using flow cytometry (FACS ARIA III) by acquiring zsGreen-positive cells. For virus production from Jurkat Tag (JTAg) cells, either 8μg NL4-3 Env-R-Luc or NL4-3 Env-R+ Luc was electroporated along with 2 μg VSV glycoprotein encoding plasmid(pMD2.G). Cells in the exponential growth phase were harvested (10^7^ cells/sample) at 300 × g for 5 min. Cells were washed using phosphate-buffered saline (PBS (1×), pH 7.0) to remove cell debris and residual serum and were resuspended in warm Opti-MEM (200μl/sample). Next, the samples (cells with plasmid) were added to a 2mm-gap electroporation cuvette (Bio-Rad, USA). The sample was pulsed at 140V and 1,000 μF with an exponential decay on a Bio-Rad GenePulserXcell module. Warm RPMI (~600 μl) with 20% fetal bovine serum (FBS) was then added to the electroporated cells, which were then transferred to a 6-well plate containing pre-warmed 10% FBS containing RPMI. After 48h, virus supernatant was collected by removing cells using centrifugation, filtered and stored for further use.

### RNA isolation, cDNA synthesis, and qPCR

For RNA isolation, cells were lysed using TRIzol (Invitrogen, Thermo Fisher Scientific, USA) and phase-separated using chloroform. The aqueous phase containing RNA was then processed using an RNA clean and concentrator kit. Samples were treated using DNaseI to remove any possible DNA contamination. RNA concentration was determined using a Qubit 4.0 fluorometer with Qubit HS RNA kit. cDNA first-strand synthesis was achieved by either random hexamers (ThermoFisher scientific, cat#SO142) or Oligo(dT) primers (cat#SO132) using Superscript III (Invitrogen, Thermo Fisher Scientific, cat#18080044 USA). For expression quantification, SYBR green-I (Invitrogen, cat#S7585) based qPCR was performed, and data were represented as log2 fold change.

### circRNA enrichment and Nanopore library preparation

For nanopore sequencing, linear RNAs were depleted by various steps to enrich circRNAs. Precisely, linear RNAs were depleted by ribodepletion, polyadenylation, and poly(A) RNA removal followed by RNaseR treatment. This method drastically depleted rRNA, poly(A) RNA, and non-polyadenylated RNAs, including highly structured RNAs, and provided a highly pure circular RNA fraction. Purified circRNA fractions were then fragmented using NEBNext Magnesium RNA fragmentation module and polyadenylated and processed for Nanopore direct-RNA sequencing protocol. A detailed description of this method is available in the supplementary information.

### Ribonuclease R treatment of the RNA

For, degradation of the linear RNA pool. 1 μg RNA was incubated with the 1 unit of RNaseR at 37°C for 30 min. RNaseR was then inactivated at 90°C for 10 minutes. RNA was then eluted using a Zymo clean and concentrator kit and used for RT-qPCR to quantify respected RNAs.

### Nanopore sequencing and Data analysis

The Direct RNA sequencing libraries were prepared according to the ONT protocol SQK-RNA002, and RNA sequencing was performed using the MinION platform with an FLO-MIN106 flow cell. The MinKNOW (v3.2.6) interface was used, and data was basecalled using Guppy (v3.2.8), keeping high accuracy basecalling. The filtered data were mapped to the human genome (hg38) using a custom command line pblat (39) software package. For capturing the circRNA from the DRS Nanopore reads, we designed a virtual backspliced junction library of all circRNA present in the circBase(40) and circAtlas(41) databases. The circRNA sequences were downloaded from circBase (hg19) and circAtlas (hg38) database, respectively. FASTA sequences were converted into backsplice FASTA sequences using the circDR-seq custom script. Then these backsplice sequences are shortened to 100bp, 50bp upstream, and 50bp downstream to the backsplice junction. This shortening was done mainly to identify the backsplice read in the Nanopore reads.

Further, using pblat, backsplice sequences were aligned to the sequencing reads (generated in this study). pblat is able to map the sequences across the reads, but when it encounters a backsplicing junction, it gets split into two segments that are individually mapped to the different regions of the same gene. Two segments of a read that match upstream and downstream of a splice site signifies a backsplice junction and, therefore, can be considered a circRNA. To qualify as a circRNA, the Blat score has to reach 60 and should be appropriately aligned 40 bp across the junction (20 upstream and 20 downstream)

### Simulation circRNA reads

To generate simulated nanopore circRNA sequencing datasets, we used an existing tool Nanosim(42). Towards this, we downloaded circBase fasta sequences and selected top 3000 circRNA sequences for generating simulated reads, and converted these 3000 circRNA linear fasta sequences into backsplice fasta sequences. Subsequently, simulated reads were generated using ‘genome mode’ provided in NanoSim. In total, 12 million reads were simulated, which resulted in aligned simulated reads fasta file and unaligned simulated reads fasta file. Only the aligned simulated reads fasta file was used for downstream analysis, consisting of back-spliced reads. Performance assessment of circDR-seq was done by precision and recall rates of circRNA identification, and the overall performance was evaluated by the F1 score using the following equation.

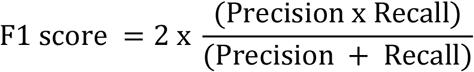

### CasRx based circRNA knockdown and infectivity assay

For circular RNA knockdown Cas13d and gRNA expressing plasmids (1:2 ratio) were co-transfected into Jurkat Tag cells (JTAg) by electroporation. Cells in the exponential growth phase were harvested (10^7^ cells/sample) at 300 × g for 5 min. Cells were washed using phosphate-buffered saline (PBS (1×), pH 7.0) to remove cell debris and residual serum and were resuspended in warm Opti-MEM (200μl/sample). 8 μg of total plasmid (2 μg of Cas13d and 4 μg gRNA expressing vectors and 2 μg pCDNA 3.1 BS(-) was then mixed into the suspended cells. Next, the samples were added to a 2mm-gap electroporation cuvette (Bio-Rad, USA). The sample was pulsed at 140V and 1,000 μF with an exponential decay on a Bio-Rad GenePulserXcell module as described previously(43). Warm RPMI (~600 μl) with 20% fetal bovine serum (FBS) was immediately added to the electroporated cells, which were then transferred to a 6-well plate containing pre-warmed 10% FBS containing RPMI. After 48h, cells were spun down at 300xg for 5 min, resuspended in fresh medium, and selected using blasticidin for four days. Cells were divided into two parts; one part was used to analyze the knockdown efficiency of the circRNAs, and the other part having the same number of cells was processed for HIV-1 infectivity analysis using HIV-1 Luc virus. For HIV-1 infectivity, luciferase-containing virus (Produced from single-cycle NL4-3 Luc (Env defected, Vpr defected, and Nef defected) was used. After 48h, cells were collected and processed for luciferase assay.

### m6A circRNA pulldown and analysis

For m6A pulldown, total RNA (10 μg) was subjected to RNaseR digestion followed by cleanup using RNA clean and concentrator kit (Zymo). Linear RNA depleted fraction was then fragmented using Magnesium RNA fragmentation module using manufacturer’s protocol. Purified RNA then was pulldown using m6A specific antibody using EpiMark N6-methyladenosine enrichment kit(NEB). RNA was converted into cDNA using random hexamer primers and Revert aid RT (Invitrogen). Selected candidates were then analyzed by RT-qPCR using specific primers. Data was normalized to input with IgG and m6A fraction.

### Luciferase assay

Luciferase assay was performed in 96-well plates to analyze viral infectivity upon circRNA knockdown. Equal numbers of cells were lysed in 96-well plates using 100 μl lysis buffer (1% Triton X-100 (Sigma Aldrich), 25 mM Tricine (VWR), 15 mM of potassium phosphate (at pH 7.8), 15 mM MgSO, 4 mM EGTA, and 1 mM DTT) for 20 minutes at room temperature. Luminescence was measured using SpectraMax-i3X plate reader (Molecular Devices, USA) by injecting 50 μl of substrate buffer (lysis buffer+ 1 mM DTT (VWR Scientific), 1 mM ATP (Sigma Aldrich), and 0.2 mM D-Luciferin (Cayman chemical, USA)) in 50 μl of cell lysate in 96-well white plates (SPL life sciences, cat #30196). Data was normalized to Bradford readings of the cell lysate in case of Viral infectivity based on VSVG pseudotyped HIV-1 Luc (NL4-3 Env-R-Luc or NL4-3 Env-R+ Luc). For LTR-Luc experiment, LTR from NL4-3 and from different transmitted founder viruses were cloned into PGL3-Luc basic vector. For analysing effect of ciTRAN knockdown on different HIV LTRs, LTR-Luc constructs along with pCasRx and pCDNA Tat were transfected into HEK293T with non-targetting gRNA(gRFP) or ciTRAN gRNA. After 48h, luciferase assay was performed to understand the effect of ciTRAN knockdown on LTR activity. Data was normalized to renilla luciferase cotransfected (10ng) in every condition. Similarly, for analysing the effect of SRSF1 and ciTRAN modulation on different LTRs, PGL3-Luc or PGL3-LTR-Luc along with vector or pCDNA Tat were tranfected in HEK293T along with SRSF1 and pCDNA 3.1(+) circMini ciTRAN. After 48 h luciferase assay was performed to analyse the effect of SRSF1and ciTRAN level on LTR ativity.

### Subcellular fractionation

Subcellular fractionation was done as described previously(44). For the cytoplasmic fraction, 1 million cells were collected at 500 × g at 4°C for 5 min. The supernatant was removed, and the cell pellet was stored on ice. Cells were resuspended gently in 380 μl ice-cold Hypotonic lysis buffer (HLB) supplemented with Ribolock (Invitrogen, Thermo Fischer Scientific, USA) RNase inhibitor. Cells were incubated for 10 min, vortexed briefly, and then centrifuged at 1000 × g for 3 min at 4°C. The supernatant containing cytoplasmic fraction was transferred to a new tube, and the pellet was stored on ice. The supernatant was mixed with 1 mL RNA precipitation solution and stored at −20°C for at least 1h. After incubation, the sample was centrifuged at 18,000 × g at 4°C for 15 min, and the pellet was processed for RNA isolation using TRIzol and DNA isolation using the phenol-chloroform method. The nuclear fraction containing the pellet was washed thrice with 1 mL of ice-cold HLB and centrifuged at 300 × g at 4°C for 2 min. Pellet was processed for DNA isolation using the phenol-chloroform method.

### Estimation of HIV-1 reverse Transcription, integration and transcription upon ciTRAN knockdown

For reverse transcription and integration analysis, DNA was isolated from the cytosolic fraction and nuclear fraction, respectively, at different time points and was analyzed using qPCR using the specific primers forward 5’-TCTGGCTAACTAGGGAACCCA-3’ and Reverse 5’-CTGACTAAAAGGGTCTGAGG −3’ in case of reverse transcription analysis. HIV-1 integration was quantified with a two-step Alu-Gag PCR where in the 1st step, genomic DNA with Alu forward 5’-GCCTCCCAAAGTGCTGGGATTACAG-3’ and Gag Reverse 5’-TACCATTTGCCCCTGGAGGTT −3’ was pre-amplified. The amplicon from this PCR was used as a template for the second round of qPCR using HIV-1 *gag*-specific primers (Forward 5’ - TCTGGCTAACTAGGGAACCCA-3’ and Reverse 5’-CTGACTAAAAGGGTCTGAGG - 3’). HIV-1 RNA abundance from cytoplasm as a measure of HIV-1 transcription was quantified at different time points using forward 5’-TTGTACTGAGAGACAGGCT −3’ and Reverse 5’-ACCTGAAGCTCTCTTCTGG-3’. RNA was isolated using TRIzol/Chloroform method and cDNA synthesis was performed using the random hexamer primer.

### Nuclear Run-on assay

A nuclear run-on assay was performed as described previously(45). For nuclei isolation, cells were harvested using ice-cold hypotonic solution (150 mM KCl (Sigma), 4 mM MgOAc (Sigma), and 10 mM Tris-HCl (Sigma), pH 7.4) and were pelleted by centrifugation 300 × g for 5 min. Next, the cell pellet was resuspended in lysis buffer (150 mM KCl, 4 mM MgOAc, 10 mM Tris-HCl, pH 7.4, and 0.5% NP-40). The nuclear run-on mixture (10 mM ATP, CTP, GTP, BrUTP, and the crude nuclei) was incubated at 28 °C for 5 min in the presence of RNase inhibitor (Invitrogen). The RNA was then isolated by TRIzol reagent (Invitrogen) as per the manufacturer’s instructions, and DNA was eliminated by DNase I (Takara) treatment. Nascent transcripts were then immunoprecipitated by anti-BrdU antibody (Merck, Cat# B8434) and converted to cDNA for qPCR analysis. qPCR analysis was done using forward 5’-TTGTACTGAGAGACAGGCT −3’ and Reverse 5’-ACCTGAAGCTCTCTTCTGG-3’ primers.

### CRISPR-assisted RNA–protein interaction detection method

We generated gRNA targeting the back-splicing junction of ciTRAN and cloned it into an empty gRNA expressing vector (Addgene #109054). The gRNA sequences were designed by keeping GC% = 40–60% and filtered for off-targets. 5 μg of gRNA-expressing plasmid (ciTRAN) and 5 μg of BASU–dCasRx constructs were cotransfected by electroporation. Before collection, Cells were washed thrice with ice-cold PBS and lysed using 500 μl lysis buffer (50 mM Tris-HCl (pH 7.4); 150 mM NaCl; 0.5% Triton X-100; 1 mM EDTA supplemented with fresh EDTA-free protease inhibitor cocktail (Roche) at 4 °C for 15 min with rotation. The lysate was spun down at 16,000 × g for 15 min at 4°C. The supernatant was quantified using Bradford (B1696, Sigma) and normalized for protein concentration. Biotinylated proteins were enriched with MyOne T1 streptavidin beads (Thermo Fisher) after 2 h of incubation at 4 °C with rotation, and 3 washes were performed with 1 ml ice-cold lysis buffer. Proteins were eluted from the beads using 2 × SDS loading buffer by incubating for 10 min at 95°C and subjected to SDS-PAGE. Following washing with clean water, silver staining was performed. The visible differential protein band was excised from the gel and processed for LC-MS analysis. Obtained spectra were run on a Mascot server to identify the target proteins.

### Immunoblotting

For immunoblotting, samples were lysed using RIPA lysis buffer and mixed with Laemmli buffer. Samples were either run on 8% or 12% tricine gels depending upon protein size. Next, gels were electroblotted on the PVDF membrane (Immobilon-FL, Merck-Millipore). The membrane was blocked using a commercial blocking buffer (BIORAD) for 5 min, followed by primary and secondary antibody incubations for one h each at room temperature. Post antibody incubation membrane was washed thrice (5 min per wash) using Tween20 containing 1× Tris-Buffered saline (TBST). Detection of beta-actin, p24, SRSF1, and GFP was carried out using anti-beta actin (LI-COR Biosciences, Cat# 926-42210, RRID: AB_1850027), mouse anti-p24 (NIH ARP), anti-SRSF1 (Merck) and anti-GFP (ABio laboratories). Details of antibodies with details is provided in supplementary resource table.

### Photoactivatable ribonucleoside-enhanced crosslinking and immunoprecipitation (PAR-CLIP)

PAR-CLIP was performed as described previously(46). E6.1 cell lines were expanded two days before the infection, and the old medium was replaced with the fresh medium before infection. Next, cells were collected and divided into two flasks, cells in one flask were challenged with HIV-1 particles at MOI 5, and the second flask was challenged with a mock referring to it as control. In the next step, the virus-containing medium is replaced after 6 h with a fresh medium. Cells were incubated with 100 mM 4-thiouridine(4-SU) for 16 h. Following incubation, cells were collected using centrifugation and washed twice with the ice-cold PBS (Hyclone). After washing, cells were resuspended in ice-cold PBS and crosslinked thrice with a one-minute gap using a 365 nm-based UVP crosslinker at 0.2 J/cm2. Following the crosslinking step, cells were collected and either stored in −80 until further use or directly processed for the pull-down. For the pull-down experiment, Protein G beads were initially incubated with the SRSF-1 (5μg) antibody or IgG at room temperature for 40 minutes. After lysing a sample with NP40 (Sigma) based lysis buffer, the lysate was cleared using centrifuge and supernatant was added to the pre-occupied antibody-beads mixture. The mixture was then incubated at 4°C overnight. Post incubation beads were washed with high-salt buffers and finally dissolved in proteinase K digestion buffer for proteinase K digestion (Thermo Fischer Scientific). After that reaction was stopped by adding 1 mL Trizol and RNA isolation and clean up was performed using Zymo clean and concentrator kit. Isolated RNA was then processed for cDNA synthesis. Similarly, PAR-CLIP was also performed for SRSF1 mutants from JTAg cells. SRSF1 mutants were cloned with HA in pCDNA3.1(-) and ectopically expressed (5μg) using electroporation. Pulldown was performed using anti-HA antibody with protein G beads and RT-qPCR analysis was performed for ciTRAN enrichment using specific primers.

### Chromatin immunoprecipitation (ChIP)

For chIP, 10 million JTAg cells per condition were cross-linked for 10 minutes at room temperature with 1% formaldehyde (Sigma) and then quenched with 0.125 M glycine for 5 minutes at room temperature(47). Cells were washed three times in cold PBS before being resuspended in a buffer containing 50 mM HEPES-KOH (pH 7.5), 140 mM NaCl, 1 mM EDTA, 10% glycerol, 0.5% NP-40, 0.25 % Triton X-100, and protease inhibitors (Thermo Scientific). Nuclei were pelleted at 800 × g for 5 minutes at 4 °C and resuspended in a buffer containing 10 mM Tris-HCl pH 8.0, 200 mM NaCl, 1 mM EDTA, 0.5 mM EGTA, and protease inhibitors, then incubated on ice for 10 minutes. Nuclei were pelleted at 800×g for 5 minutes at 4 °C and resuspended in a buffer containing 10 mM Tris-HCl pH 8.0, 200 mM NaCl, 1 mM EDTA, 0.5 mM EGTA, and protease inhibitors, then incubated on ice for 10 minutes. The nuclei were collected and resuspended in a sonication buffer containing 10 mM Tri-HCl (pH 8.0), 100 mM NaCl, 1 mM EDTA, 0.5% EGTA, 0.1 % Sodium deoxycholate, and 0.5 percent N-lauryl sarcosine, as well as protease inhibitors.

DNA was sonicated using a probe sonicator for 15 cycles of 30 seconds and 45 seconds off to obtain an average fragment length of 300–700 bp. Samples were centrifuged at 16000xg for 10 minutes at 4 °C after being treated with 1% Triton X-100. During each ChIP cycle, an aliquot of sonicated DNA was reverse-crosslinked and run on a 1% agarose gel to confirm fragment size.

Chromatin (25 μg) from different samples was immunoprecipitated by adding the antibody of interest (anti-SRSF1, anti-RNAPII and anti-Tat), followed by overnight incubation at 4 °C. After overnight incubation, 30 μl of Protein G beads (Invitrogen) were added and incubated for 1 h at 4 °C. Beads were washed sequentially for 3 min each in low salt (20 mM Tris-HCl pH 8.0, 150 mM NaCl, 2 mM EDTA, 0.1% SDS, 1% Triton X-100), high salt (20 mM Tris-HCl pH 8.0, 500 mM NaCl, 2 mM EDTA, 0.1% SDS, 1% Triton X-100), LiCl buffer (10 mM Tris-HCl pH 8.0, 0.25 M LiCl, 1% NP40, 1% Sodium deoxycholate) and TE buffer. Beads were eluted in 150 μl elution buffer (50 mM Tris-HCl pH 8.0, 10 mM EDTA, 1% SDS, 50 mM NaHCO3) and treated with 1 μl RNaseA (1mg ml^-1^ Ambion) at 37 °C for 30 min. Cross-linking was reversed, and proteins were degraded by adding 1 μl proteinase K and incubation at 65°C for 4 h. Eluted DNA was purified and used for qPCR analysis. For ChIP experiment, four different conditions were used. TAR enrichment for SRSF1, RNAPII and HIV-1 Tat ChIP was checked in non-targetting gRNA expressing JTAg cells (gRFP, termed as vector), ciTRAN knockdown cells (gciTRAN), ectopic expression of SRSF-1 highlighted with underline (pCDNA3.1 (-)) along with pCDNA 3.1(+) circMini empty vector and pCDNA 3.1(+) circMini ciTRAN. Data was represented as fold change for TAR enrichment normalized with IgG background.

### Virion incorporation of Vpr

Virus incorporation assay was done as described previously(43). Virus particles (HIV-1 Luc) were produced by transfecting HEK293T cells using calcium phosphate transfection reagent in a 10^cm2^ plate, NL4-3 Env-R-Luc, pMD2.G (VSVG) along with 2μg Vpr expressing construct (pEGFP-C2-Vpr) or vector alone. After 12-15 h post-transfection, the medium was replaced with 2% FBS containing DMEM. After 48h, the virus-containing supernatant was collected at 500 x g for 5 min to exclude any cell debris. Next, the virus-containing supernatant was filtered using a 0.22-μm syringe filter (Whatman). The suspension was overlaid on a 20% sucrose (prepared in 1 × PBS) cushion and concentrated at ~110,000 × g for 2h at 4°C using a Beckman-Coulter ultracentrifuge. After the spin, the supernatant was discarded, and the pellet was either suspended in 1xPBS for infectivity experiment or in Laemmli buffer containing 50 mM Tris(2-carboxyethyl) phosphine hydrochloride (TCEP) for western blot.

### SRSF1 deletion mutants

SRSF-1 full-length gene was cloned in pCDNA 3.1(-)HA vector using XbaI and BspEI after PCR amplification from cDNA using specific primers. Different mutants of SRSF-1 such as RRM1, RRM2, and RS, were then generated on this backbone, keeping HA in the frame using PCR mutagenesis (primers are provided in the supplementary table S1). Cloning was confirmed by Sanger sequencing, and the expression of each construct was confirmed using anti-HA immunoblotting. Further, the CPP-tagged SRSF1 RRM2 mimic was cloned in house. The RRM2 domain was cloned into an SP-CPP sequence containing plasmid which was previously generated in the lab. The CPP sequence used is GAYARKAARQARAGVD. The CPP-tagged RRM2 has CMV promoter and HA tag at the C-terminus facilitating its purification using anti-HA antibody-based pull-down in mammalian cell culture supernatant. Since the CPP-RRM2 plasmid generated in this study contains a CMV promoter for mammalian expression therefore the experiment was performed in transfection format by cotransfecting SP-CPP or SP-CPP-RRM2-HA along with HIV-1 Tat and PGL3-LTR-Luc constructs. Luciferase readings as a function of LTR activity was analyzed to confirm the functionality of the construct.

### PBMC isolation from peripheral blood

Blood was collected in preservative-free anticoagulant (0.2 % Final concentration of EDTA) from three donors and processed immediately for PBMC isolation. PBMC isolation was done with the help of Histopaque using the manufacturer’s protocol (Sigma Aldrich Cat No. 10771). Precisely, 3 ML anticoagulated blood was layered on the top of an equal amount of Histopaque-1077. Afterward, the sample was carefully centrifuged in a swing bucket rotor keeping acceleration and brake at the lowest setting at 400x g for 30 minutes at room temperature. Following centrifugation, cells from the opaque interface were transferred to a fresh centrifuge tube. Cells were washed twice with isotonic phosphate buffer saline collected upon centrifugation at 250 × g for 10 minutes. Finally, cells were re-suspended in the isotonic phosphate buffer saline or RPMI 1640 (BIOWEST) and processed further for the CD4+ cell isolation.

### CD4^+^ cell purification

CD3^+^/CD4^+^ T cells were purified by positive selection using magnetic separation with a CD4^+^isolation kit (Miltenyi Biotec) using PBMCs from three donors as per the manufacturer’s protocol. Purified CD4^+^ T cells were characterized following counter-staining with anti-CD4-APC antibody (1:200) using flow-cytometry (Miltenyi Biotec), and anti-CD3-FITC labelled antibody. Antibodies were diluted in PBS / 1% BSA / 0.05% NaN_3_ (PBA).

### Primary T cells maintenance

Primary CD4+ T cells were grown and maintained in RPMI-1640 (Biowest) supplemented with 10% FBS. For expanding the CD4^+^ T cells, the cells were activated using 5 μg/mL phytohaemagglutinin (PHA, Sigma-Aldrich) and 50 IU/mL recombinant human IL-2 (Gibco). IL-2 was also supplemented to each culture for the maintenance of the cells.

### Statistical analysis

Statistical analyses were performed using GraphPad Prism 9.0. Data were presented as the means ± standard deviation (SD). The two-tailed Student’s t-test (paired/unpaired) or one-way ANOVA (multiple comparisons) was used to assess the significance between two or more groups. All reported differences were *p < 0.05, **p < 0.01, ***p < 0.001 and ****p < 0.0001 unless otherwise stated.

## Supplemental figures

**Figure S1.**
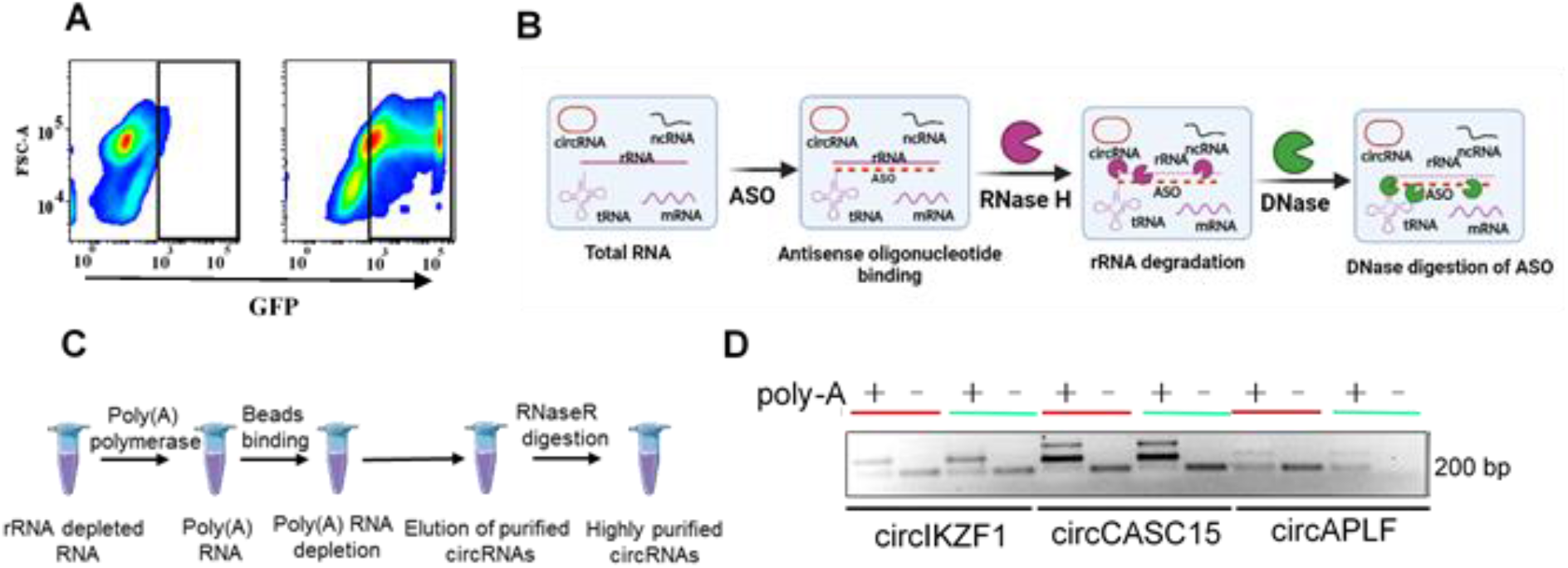
Procedures and checks for circRNA enrichment from the infected and mock Jurkat T cells. **(A)** Flow-cytometry of mock and HIV-1 GFP infected E6.1 T cells. **(B)** Schematics depicting sequential steps for depleting rRNAs by recruting ASO followed by RNAseH mediated degradation (Refer to supplementary information for more details). **(C)** Schematics depicting sequential steps to deplete linear RNAs by polyadenylation, oligo(dT) magnetic bead-based removal, and RNaseR treatment to enrich circRNA fraction. **(D)** Validation of circRNA fragmentation and addition of polyA by oligo(dT) primed cDNA synthesis and PCR in mock(red) and infected(green) samples.

**Figure S2.**
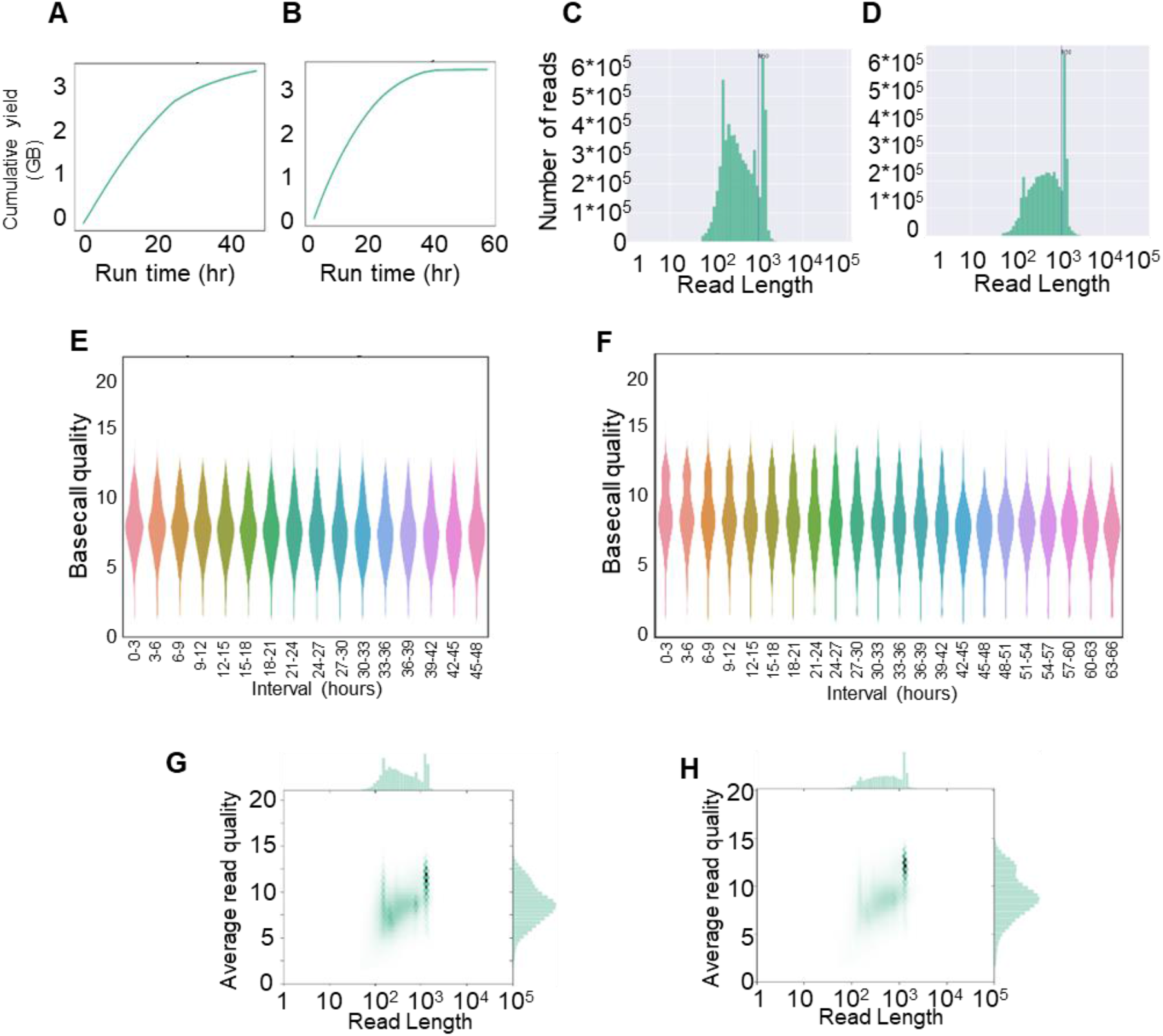
Quality checks for Nanopore library and sequencing. **(A,B)** the representative cumulative yield is shown as a parameter of flow cell performance over time in mock-treated (**A**) and infected (**B**) cells. **(C,D)** Read-length distribution analysis of mock-treated (**C**) and infected cells (**D**). **(E,F)** qualitative analysis of basecalled sequencing reads over time obtained from mock-treated (E) and Infected cells (**F**). **(G,H)** qualitative analysis of read length and read quality of sequencing reads obtained from mock-treated (**G**) and Infected cells (**H**).

**Figure S3.**
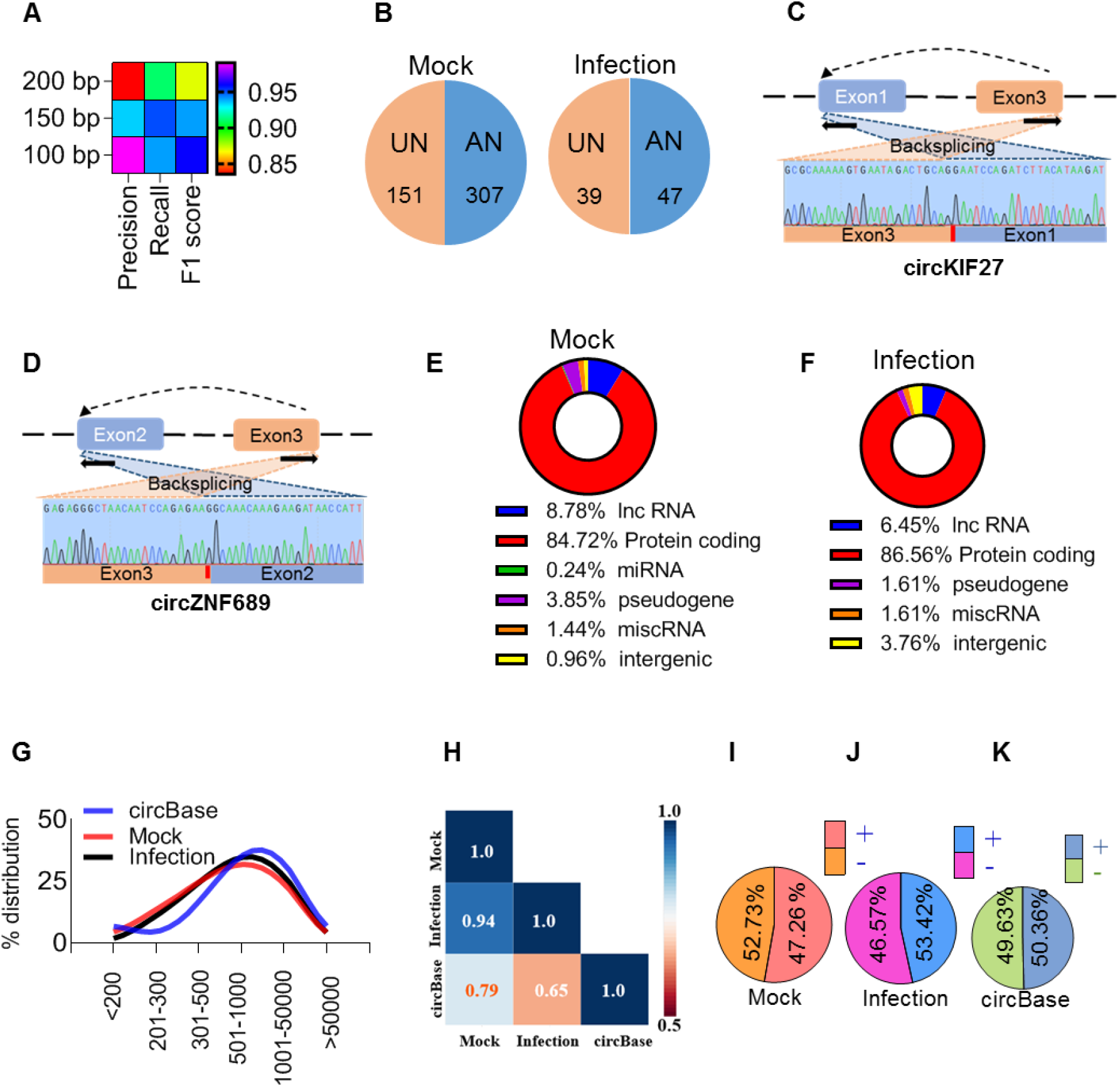
Nanopore data analysis, pipeline validation and identification of novel circRNAs. **(A)** Qualitative analysis of various libraries using precision, recall, and F1 score after simulation. **(B)** Unannotated (UN) and annotated (AN) circRNAs in mock and infection. **(C,D)** Validation of unannotated circRNAs (from **B**) for the presence of novel back-spliced junction by Sanger sequencing. **(E,F)** cataloguing of circRNAs according to the RNA classes **(G)** circRNAs length distribution obtained herein and its comparison with those reported in circBase. **(H)** circRNA length-distribution correlation in mock infection with circBase database. (**I,J,K)** Strand assignment of detected circRNAs (**I,J**) and its comparison with the circBase database for occurrence (**K**).

**Figure S4.**
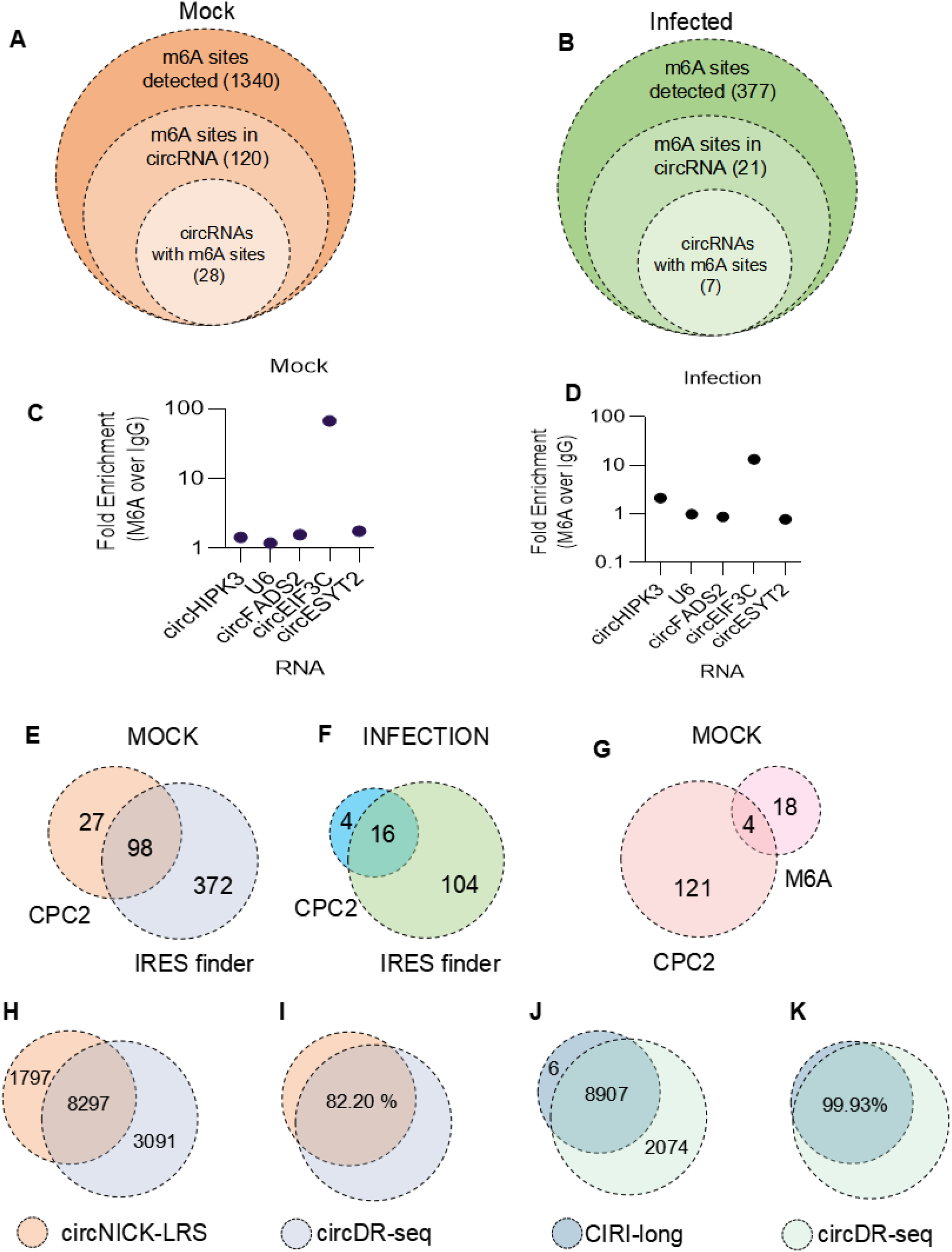
Detection of m6A, protein-coding potential of circRNAs, and comparison of circDR-seq with published pipelines. (A,B) Detection of M6A among the reads obtained and M6A presence in the circRNAs. (C,D) m6A profile of randomly selected circRNAs from Mock and infection revealed by m6A pulldown followed by RT-qPCR. (E,F) The protein-coding potential of circRNAs assumed by IRES presence. (G) An overlap between M6A modified circRNA with circRNAs having coding potential as estimated using CPC2. (H,I) A comparison of circDRseq with circNICK-LRS and the percentage overlap between circRNA estimation from the data generated by the circNICK-LRS paper. (J,K), A comparison of circDRseq with CIRI-long and the percentage overlap between circRNA estimation from the data generated by the CIRI-long paper.

**Figure S5.**
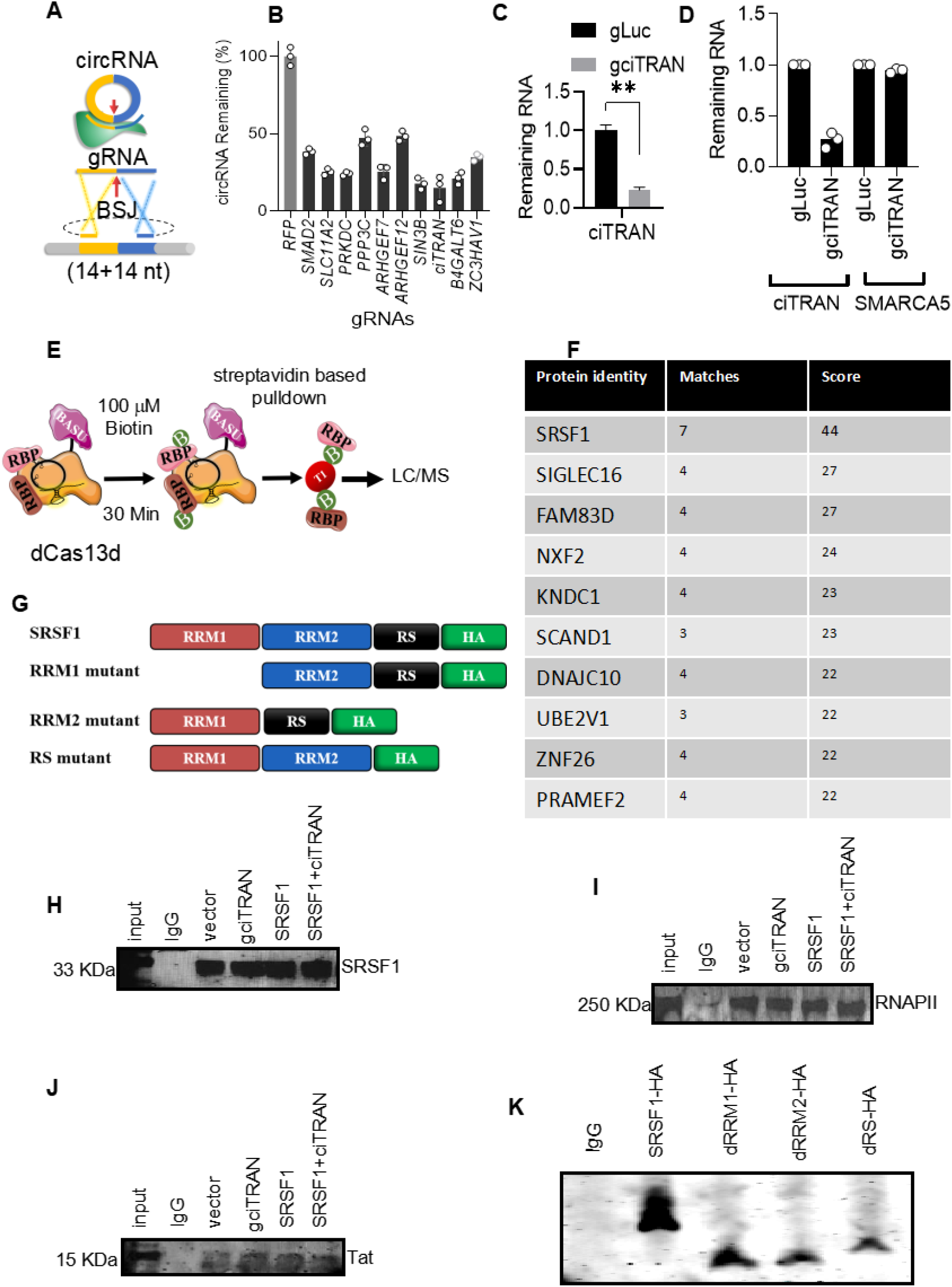
CRISPR, proximity-labeling proteomics, mutant analysis, chromatin immunoprecipitation, and PAR-CLIP-associated checks. **(A)** gRNA design schematics for a circRNA knockdown by CasRx. **(B)** Levels of ciTRAN after CRISPR/CasRx knockdown. **(C)** Levels of ciTRAN after knockdown (for Figure-3). **(D)** qRT-PCR analysis of ciTRAN and its linear counterpart SMARCA5 RNA (parental gene encoding ciTRAN) upon ciTRAN knockdown. **(E)** Schematics showing CasRx-BASU-based proximity ligation. **(F)** Proteins identified after LC/MS analysis. **(G)** Deletion mutants tagged with HA of SRSF1 generated by PCR mutagenesis. (**H-J)** Immunoblotting after chromatin immunoprecipitation (ChiP) of SRSF1(**H**), RNAPII (**I**) and HIV-1 Tat (**J**) using specific antibodies. **(K)** Immunoblot for SRSF-1 mutants using anti-HA antibody after PAR-CLIP.

**Figure S6.**
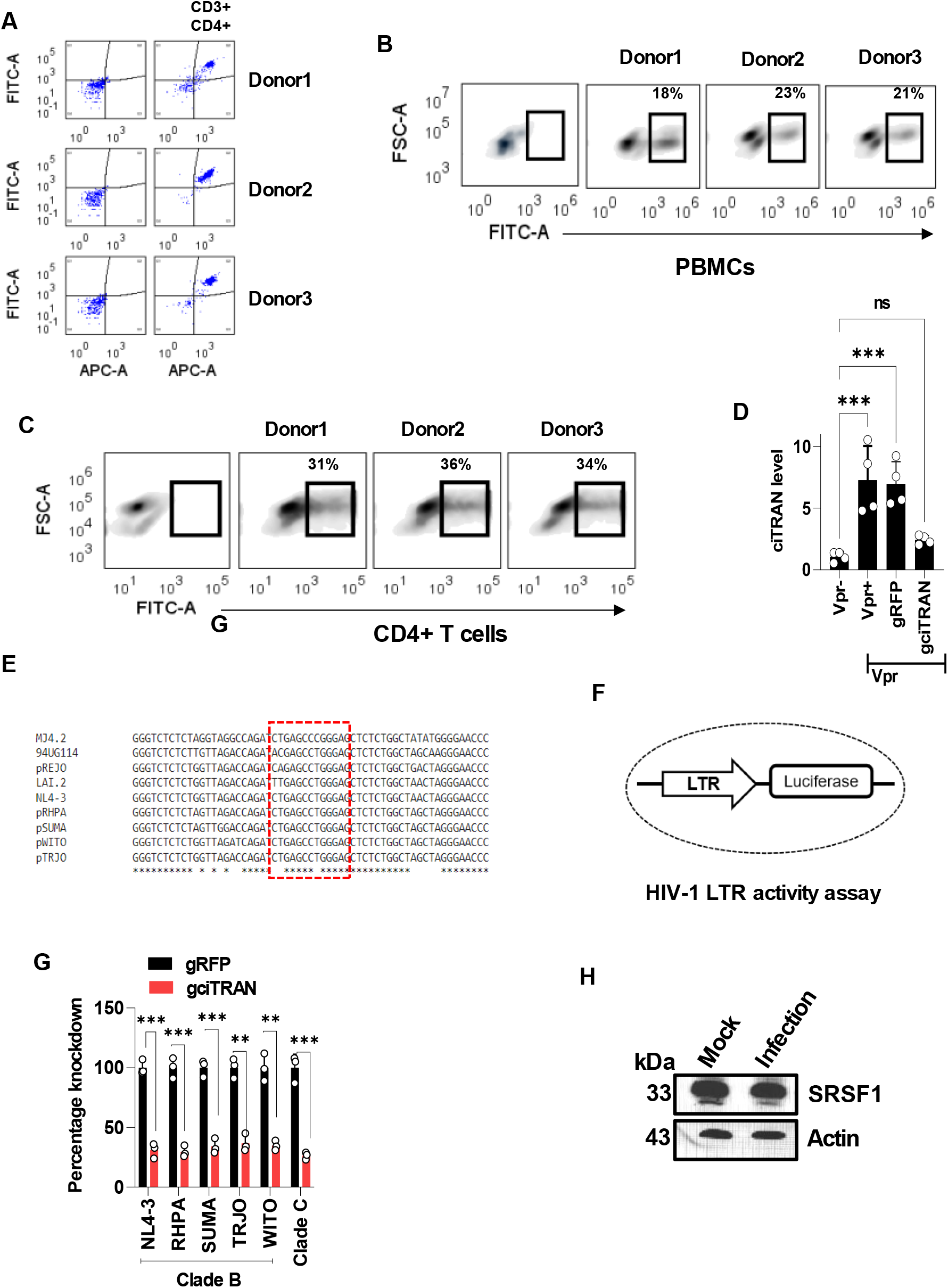
Vpr association, primary cell purification, infectivity and conservation across different clades. (**A**) Flow cytometry of CD3+/CD4+ cells purified from PBMCs of three donors. (**B,C**) Infectivity in PBMCs (**B**) and primary CD4+ T cells (**C**) from 3 Donors infected using VSVG pseudotyped NLBN zsGreen at MOI 5 for 48 h. (**D**) ciTRAN level induced by Vpr across different conditions and CRISPR-mediated knockdown validation. (**E**) SRSF-1 binding site in the LTRs across different clades and transmitted founder viruses. (**F**) LTR-luciferase reporter minigene construct(PGL3 backbone). (**G**) knockdown of ciTRAN using Cas13d in various conditions. (**H**) Effects of infection on SRSF-1 expression levels. Actin served as a loading control.

## Notes

### Competing Interest Statement

The authors have declared no competing interest.

